# Barcode 100K Specimens: In a Single Nanopore Run

**DOI:** 10.1101/2023.11.29.569282

**Authors:** Paul DN Hebert, Robin Floyd, Saeideh Jafarpour, Sean WJ Prosser

## Abstract

It is a global priority to better manage the biosphere, but action needs to be informed by monitoring shifts in the abundance and distribution of species across the domains of life. The acquisition of such information is currently constrained by the limited knowledge of biodiversity. Among the 20 million or more species of eukaryotes, just a tenth have scientific names. DNA barcoding can speed the registration of unknown animal species, the most diverse kingdom of eukaryotes, as the BIN system automates their recognition. However, inexpensive analytical protocols are critical as the census of all animal species will require processing a billion or more specimens. Barcoding involves DNA extraction followed by PCR and sequencing with the last step dominating costs until 2017. By recovering barcodes from highly multiplexed samples, the Sequel platforms from Pacific BioSciences slashed costs by 90%, but these instruments are only deployed in core facilities because of their expense. Sequencers from Oxford Nanopore Technologies provide an escape from high capital and service costs, but their low sequence fidelity has, until now, kept analytical cost above Sequel. However, the improved performance of its latest flow cells (R10.4.1) might erase this differential. This study demonstrates that a regular MinION flow cell can characterize an amplicon pool derived from 100,000 specimens while a Flongle flow cell can process one derived from several thousand. At $0.01 per specimen, DNA sequencing is now the least expensive step in the barcode workflow. By coupling simplified protocols for DNA extraction with ultra-low volume PCRs, it will be possible to move from specimen to DNA barcode for $0.10, a price point that will enable the census of all species within two decades.

## INTRODUCTION

The past 20 years have shown the effectiveness of DNA barcoding for both specimen identification and species discovery (Hebert et al. 2003, Nagy et al. 2013, Seifert. 2009, Von Cräutlein et al. 2011, Antil et al. 2023, Chac and Thinh 2023). Advances in high-throughput sequencing over this interval have enabled new approaches (eDNA, metabarcoding) which infer the species that contributed to DNA extracts prepared from environmental samples or bulk collections of specimens (Valentini et al. 2016, Hallam et al. 2021, Basset et al. 2022, Cote et al. 2023). However, these workflows require access to a reference library of barcode sequences constructed through the analysis of single specimens. Reflecting a generational effort, the DNA barcode reference library now provides coverage for 1.2 million species, 80% from the animal kingdom. Growth of the BOLD library (Ratnasingham and Hebert 2007) accelerated as analytical costs were reduced by using liquid-handlers to support DNA extraction (Ivanova et al. 2006) and to reduce PCR volumes (Hebert et al. 2013) and, most importantly, by the shift from Sanger sequencing. The transition to SMRT analysis (Rhoads *et al*. 2015) on Sequel reduced sequencing from $7.50 to $0.25 per specimen because it could characterize amplicon pools derived from ten thousand specimens (Hebert et al. 2018). By analyzing 5-fold larger pools, Sequel II lowered sequencing to $0.05 per specimen. While Sequel platforms generate high fidelity barcodes inexpensively, they have two limitations. First, their high capital ($500K) and annual service ($50K) costs restrict deployment to core facilities. Second, their flow cells are sufficiently expensive ($2K) to impede protocol development. Oxford Nanopore Technologies (ONT) removes these barriers as its base sequencer ($1000) and flow cell ($80) are both inexpensive.

Nanopore sequencing captures the nucleotide array of individual DNA molecules as they pass through nanopores (MacKenzie and Argyropoulos 2023). Within a decade of its first demonstration (Kasianowicz et al. 1996), the transformative potential of nanopore sequencing was recognized (Branton et al. 2008). Six years later, ONT released the first sequencers and flow cells to use this technology, but their high error rates constrained adoption (Laver et al. 2014). Since then, ONT has advanced sequence fidelity by improved base-calling software, by new chemistry (20+), by slowing the translocation of DNA through nanopores, and by shifting from one to two constriction points where the nucleotides are read (van der Verren et al. 2020).

The earliest applications of ONT for biodiversity science focused on gathering sequence data in remote settings (Pennisi 2017). For example, participants in the 2017 iBOL conference in Kruger National Park (South Africa) used ONT to obtain DNA barcodes from specimens during the meeting. Subsequent work demonstrated the value of ONT in supporting DNA barcode analysis in lab settings (Srivathsan et al. 2018, Loit *et* al. 2019, Srivathsan et al. 2021, Cuber et al. 2023), but relatively high error rates meant that many (100+) reads were required to generate a high-fidelity consensus sequence, keeping costs well above those achieved with Sequel. Because the latest ONT flow cells (R10.4.1) generate sequences with fewer indels and base substitutions (Ying et al. 2023, Zhang et al. 2023, Srivathsan et al. 2023), we tested their capacity to recover DNA barcodes from highly multiplexed amplicon pools (2,280–100,320 specimens).

## MATERIALS AND METHODS

### Specimens

Our work examined barcode recovery from 100,320 Malaise-trapped arthropods that were held at −20°C in 95% ethanol until processed (**Supplemental Table 1**). Prior to DNA extraction, each specimen small enough to fit in the well of a 96-well plate was imaged using a Keyence VHX-7000 digital microscope while a DSLR macro system was used to photograph larger specimens. Visual inspection enabled the placement of each specimen to class, order, and family. This analysis showed that 99.6% of the specimens were insects while the rest were arachnids or collembolans. Species in five insect orders dominated the assemblage: Diptera–76%, Hymenoptera–9%, Coleoptera–5%, Lepidoptera–4%, Hemiptera–4% with the other 1.6% belonging to 15 small orders. Overall, the 100,320 specimens included representatives of 377 families and 21,635 unique BINs, a species proxy (Ratnasingham and Hebert 2013).

### Molecular Approach

Our work evaluated the capacity of ONT to generate high-quality DNA barcode records. The three flow cells produced by ONT have 85-fold variation in pore counts (Flongle–126, MinION–2048, PromethION– 10,700) and 290-fold variation in sequence generation (1 GB, 50 GB, 290 GB) reflecting their differing pore counts and run times (1 day for Flongle, 3 days for MinION/PromethION). Because PromethION flow cells are not compatible with ONT’s least expensive sequencer (MinION), we only considered the Flongle and MinION flow cells. Before proceeding, we evaluated performance aspects of both flow cells, especially Flongle, because it is a beta product. As ONT warranties that its flow cells possess a minimum pore count that is roughly 40% of the design criterion (e.g., Flongle = > 50 pores), we evaluated the incidence of non-compliance. We also considered other factors that might impact sequence recovery and hence experimental design. ONT only warranties Flongles for 30 days so we evaluated if their performance deteriorated during this window or if they remained functional beyond the warranty period. We also evaluated the extent of variation in read count among flow cells, the factors that influence it, and read fidelity. The resulting information aided selection of the number/type of flow cells needed to assess barcode recovery at three levels of multiplexing (2K, 10K, 100K). Because it is impossible to generate identical concentrations of amplicons across a large set of PCR reactions, success in sequence recovery from pooled samples is influenced by read depth. To aid recovery, we set a target of 100 base-called reads per specimen meaning > 200K reads for the 2K pool, > 1M reads for the 10K pool, and > 10M reads for the 100K pool.

The following text provides an overview of molecular protocols.

#### DNA Extraction

DNA extracts were made from intact small specimens or from single legs of larger specimens using SPRI, solid-phase reversible immobilization (Hebert et al. 2018) and stored at −20°C until use. All extractions were performed in 96-well plates with well H12 reserved as a negative control.

#### PCR

The COI barcode region was amplified using a standard primer set with indexing protocols varying with the depth of multiplexing. PCR employed Platinum Taq (Thermo Scientific) together with standard COI barcoding primers (C_LepFolF/C_LepFolR) (Hebert et al. 2018) that were asymmetrically indexed with UMIs, Unique Molecular Identifiers (**Table 1**). For the 2K experiment, reaction components consisted of 6.25 µl of 10% trehalose (Fluka Analytical), 1.62 µl of molecular grade water (Hyclone), 1.25 µl of 10X PlatinumTaq buffer (Thermo Scientific), 0.625 µl of 50 mM MgCl2 (Thermo Scientific), 0.0625 µl of 10 mM dNTPs (KAPA Biosystems), 0.625 µl of 2 μM forward primer (IDT), 0.0125 µl of 100 μM reverse primer (IDT), 0.06 µl of 5 U/μl PlatinumTaq (Thermo Scientific), and 2 µl of DNA for a total reaction volume of 12.5 µl. For the 10K and 100K experiments, reaction volumes were cut in half to reduce analytical costs. All PCRs employed 45 cycles of amplification and standard thermocycling regime (94°C for 2 min; 5 cycles of [94°C for 40 sec, 45°C for 40 sec, 72°C for 1 min]; 40 cycles of [94°C for 40 sec, 51°C for 40 sec, 72°C for 1 min]; 72°C for 2 min).

**Table 1:**
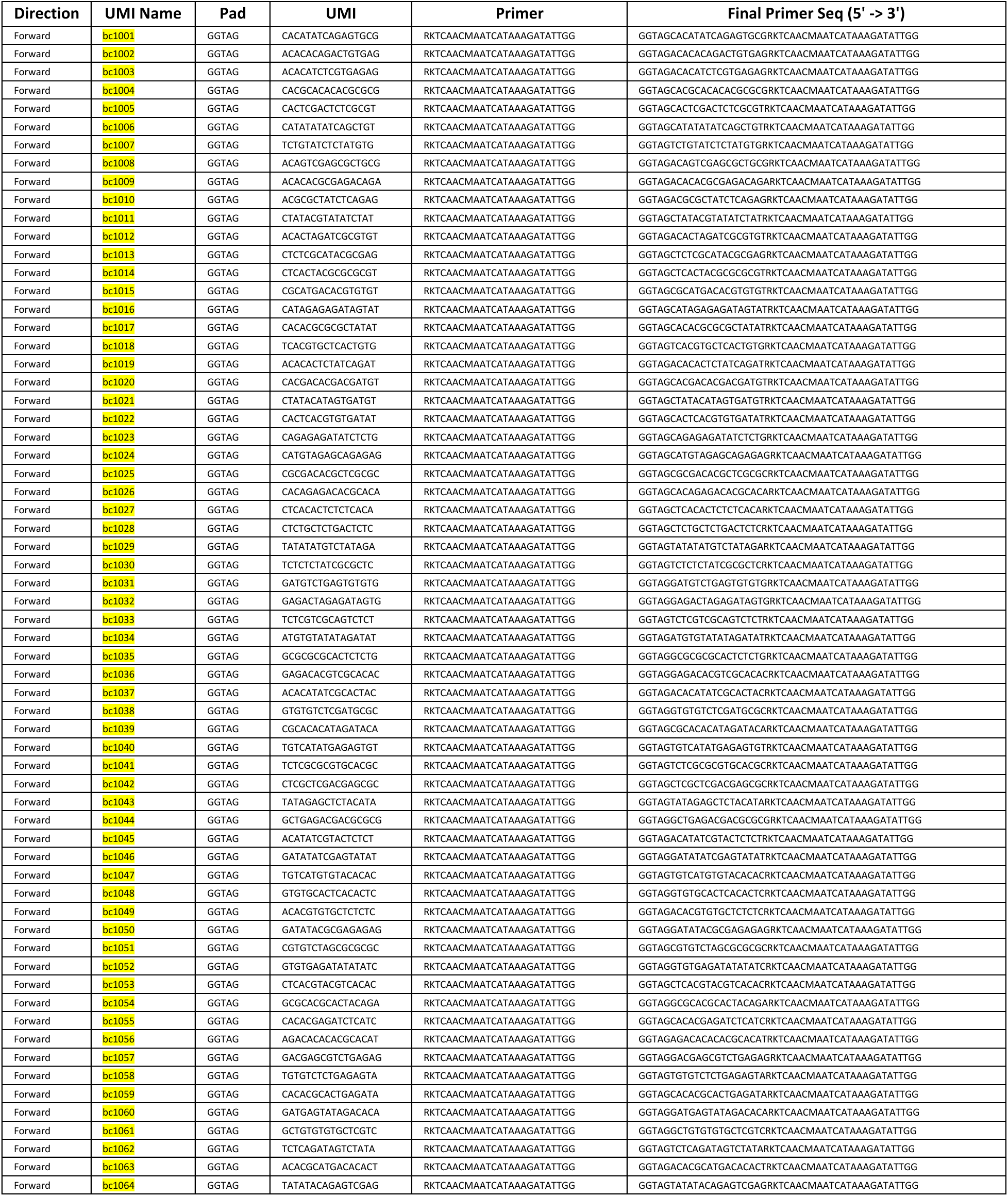
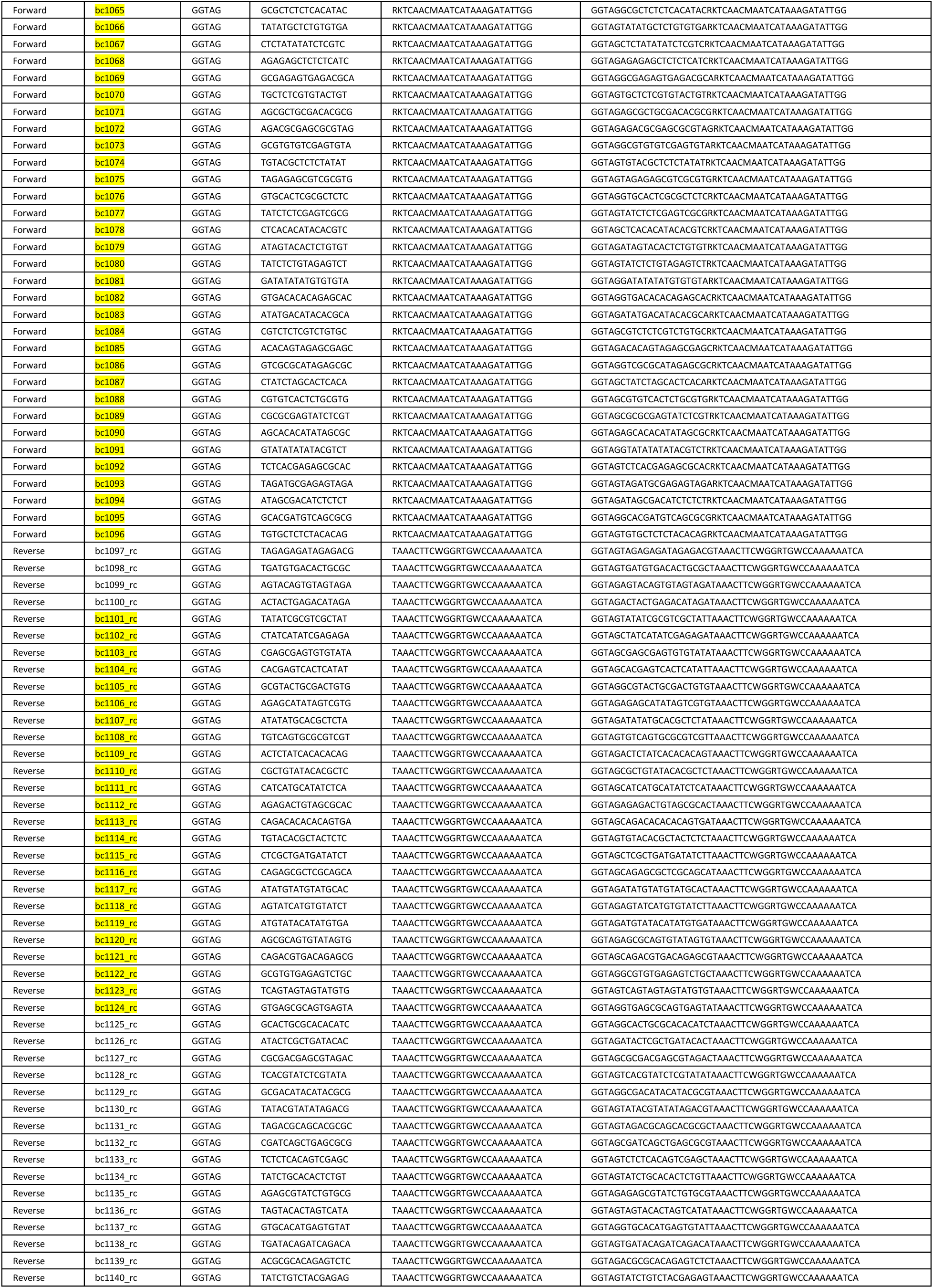
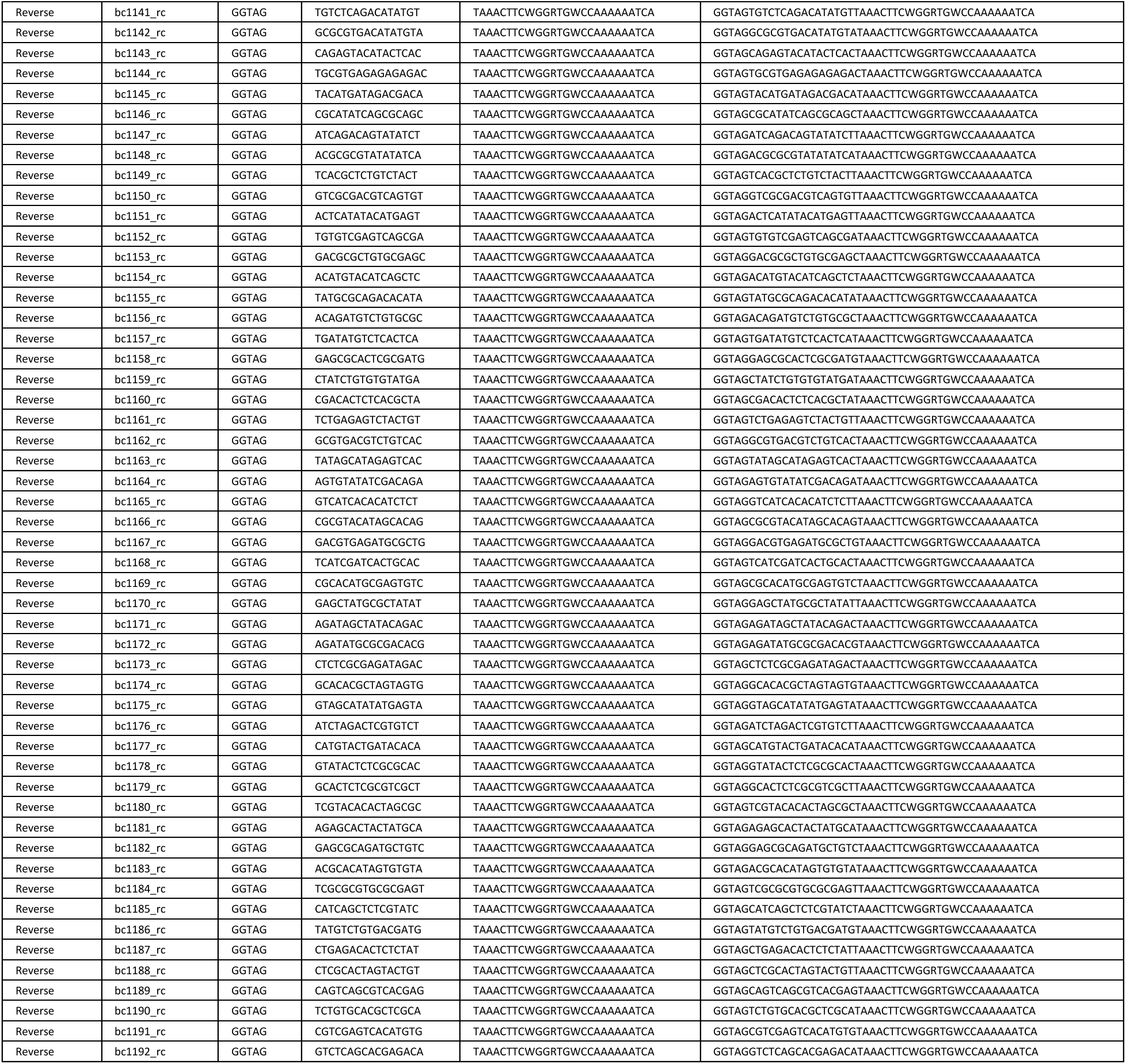
Details of the 96 forward and 96 reverse primers used in this study. The final primer sequence is composed of a pad (which shields the UMI from end-degradation), a UMI, and a COI primer. The UMIs highlighted in yellow were used for the 2K experiment (96 forward + 24 reverse), while all primers were used for the 10K experiment (96 forward + 96 reverse) and 100K experiment (96 forward + 96 reverse + 11 ONT native barcodes).

#### Multiplexing

Sequence recovery was assessed at three levels of amplicon pooling (2K, 10K, 100K). The 2K pool included amplicons from 24 96-well plates while the 10K pool included amplicons from 96 96-well plates. Finally, the 100K pool included amplicons from 1,056 96-well plates.

To attribute each sequence to its source specimen, the amplicons from each DNA extract were asymmetrically tagged with a unique combination of UMIs (**Table 1**). For all three pools, one UMI was incorporated into the forward primer (hereafter “forward UMIs”) while the other was incorporated into the reverse primer (hereafter “reverse UMIs”). Because of the 50-fold variation in multiplexing, a slightly different strategy was necessary for each pool. The 2K pool utilized 96 forward UMIs and 24 reverse UMIs, a different one for each plate, to produce 2,304 unique combinations (2,280 samples + 24 negative controls). The 10K pool required 96 forward UMIs and 96 reverse UMIs to produce 9,216 unique combinations (9,120 samples + 96 negative controls). The 100K pool utilized the same 9,216 unique UMI combinations to generate 11 amplicon pools that were subsequently differentiated by using 11 ONT native barcodes (SQK-NBD114.24). This resulted in 101,376 unique combinations (100,320 samples + 1,056 negative controls).

#### SMRT Sequencing

Prior to ONT analysis, all specimens were sequenced in batches of 9,120 on a Sequel or Sequel II platform using a standard SMRT sequencing protocol with a CCS of 99.9% (Hebert et al. 2018). This work generated a COI sequence for 91,172 of the 100,320 specimens (recovery failed in 8.8%).

#### ONT Sequencing

All sequence analysis used R10.4.1 flow cells, Q20+ chemistry, and the latest ONT base-calling software. UMI indexing was done during PCR so ONT library preparation was performed on a single pool of amplicons for each level of multiplexing, following manufacturers’ recommendations for R.10.4.1 sequencing. Briefly, amplicons were prepared for sequencing adapter ligation by end prep (NEBNext® Ultra™ II End Repair/dA-Tailing Module - E7546L), followed by purification using AMPure-XP beads. For the 100K pool, ONT Native Barcodes (SQK-NBD114.24) were ligated at this point and products purified with AMPure-XP beads before pooling. ONT sequencing adapters (SQK-LSK114) were then ligated onto the amplicons (NEB Quick T4 DNA Ligase) and purified with AMPure-XP beads. The 2K and 10K libraries were loaded onto a Flongle (FLO-FLG114) while the 100K library was analyzed on a MinION (FLO-MIN114).

### Bioinformatics

Sequence analysis was performed on an Alienware Aurora R13 running Ubuntu 22.04.3 LTS. It was equipped with an Intel Core i9 (24 core) processor, 32 GB of RAM, a GeForce RTX 3080 GPU with 10 GB RAM, and a 1 TB SSD hard drive. A custom bash script was created using freely available tools. First, the raw nanopore data (.pod5 files) was base-called using Dorado v0.3.1 and model dna_r10.4.1_e8.2_400bps_sup@v4.2.0 (https://github.com/nanoporetech/dorado). The raw reads were then filtered for length (550–950 bp) and demultiplexed using Dorado for native barcodes and Cutadapt v4.4 (Martin 2011) for the UMIs. Reads lacking a recognizable UMI (after allowing up to two mismatches) at both ends were discarded. Primer sequences were trimmed using Cutadapt and reads were discarded if they lacked a recognizable primer (after allowing up to two mismatches) at either end or if they fell below a minimum size (550 bp) after removal of the primers. The remaining reads were clustered into contigs using VSEARCH v2.21.1 (Rognes et al. 2016) with a threshold of 95%. Contigs with fewer than five reads were discarded and a majority consensus sequence was generated for each remaining contig using the Biostrings (Pagès 2022), MUSCLE (Edgar 2004) and DECIPHER (Wright 2016) packages in R (R Core Team, 2022). To mitigate contig over-splitting arising from polymerase and sequencing errors, the initial contig consensus sequences were clustered using VSEARCH with an OTU threshold of 98%. A consensus sequence was generated for each new contig and the taxonomy of its source organism was assigned using the VSEARCH SINTAX algorithm with a minimum probability threshold of 60%. The resultant sequences were filtered based on the correspondence between the ordinal taxonomy for each source specimen made by SINTAX and that based on inspection of its photograph. When a specimen possessed two or more sequences that matched the taxonomy of the source specimen, the one with the highest read depth was retained. Finally, to remove any bases outside the target, sequences were aligned in batches of 100 and their ends were clipped to the exact start and end sites of the barcode region. Base calling and subsequent data analysis required 1.5h for the 2K pool (0.75h for base calling, 0.75h for data analysis), 3.25 hours for the 10K pool (0.75h for base calling, 2.5h for data analysis), and 40h for the 100K pool (20h for base calling, 20h for data analysis).

When both platforms recovered a sequence from a particular specimen, a custom python script (pairwise_divergence.py) was used to compare them. Each pair of sequences was aligned using the Bio.Align.PairwiseAligner function from Biopython (with parameters match_score=1.0, open_gap_score = −10, extend_gap_score = −1), and the number of non-identical positions was determined (only A/G/C/T were counted; gaps, Ns or other IUPAC codes were treated as missing data and ignored). The number of non-identical bases was then divided by the total number of positions to calculate a pairwise percent divergence.

Because the ONT sequences in this study were generated automatically, the success in sequence recovery is conservative. Manual curation could elevate recovery in two ways: 1) by rescuing sequences excluded from analysis because they possessed indels linked to homopolymer tracts in the COI gene of their source specimen and 2) by allowing the inclusion of specimens with 3-4 very similar or identical sequences.

All bioinformatics scripts are available in **Supplemental File 1**. All data (images, collection details, taxonomic assignments, sequences) are available in six datasets on BOLD. The ONT records are in DS-ONT2K, DS-ONT10K, and DS-ONT100K while the Sequel records are in DS-SQL2K, DS-SQL10K, and DS-SQL100K.

## RESULTS

### PERFORMANCE OF FLOW CELLS

Among 100 Flongles, 10% failed to meet the warrantied pore count. The others generated an average of 451,000 reads but showed 40-fold variation (**Figure 1**). This variation in read count was not linked to age of the Flongle as performance remained stable for 150 days following delivery, five times the warrantied expiration date (**Figure 1**).

**Figure 1:**
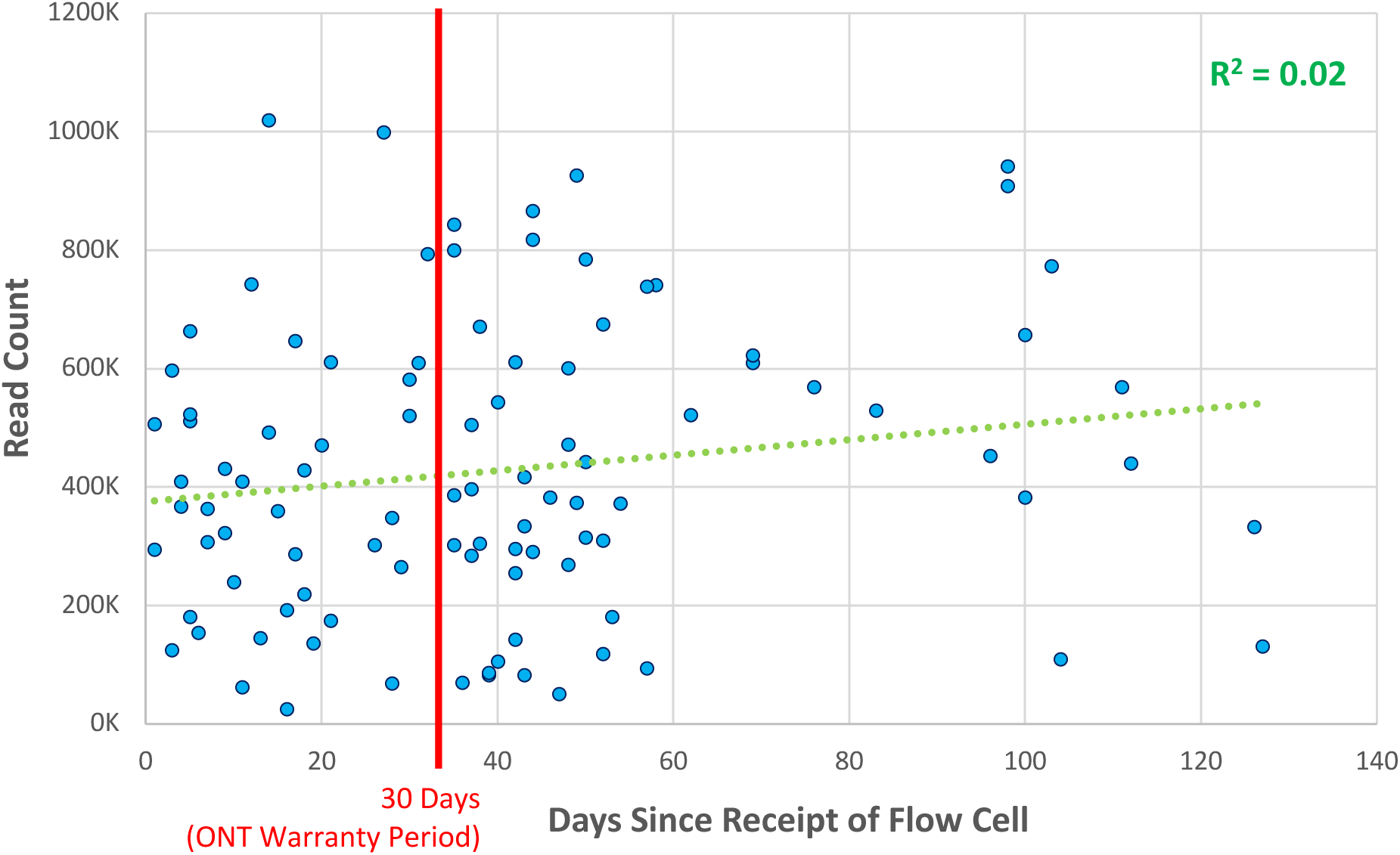
Relationship between the age of a Flongle flow cell and the number of reads it generated.

Further examination of the variation in read counts revealed a weak positive association (r^2^ = 0.28) with pore count (**Figure 2**), but the number of reads generated by a Flongle after four hours was a strong predictor (r^2^ = 0.81) of its final productivity (**Figure 3**).

**Figure 2:**
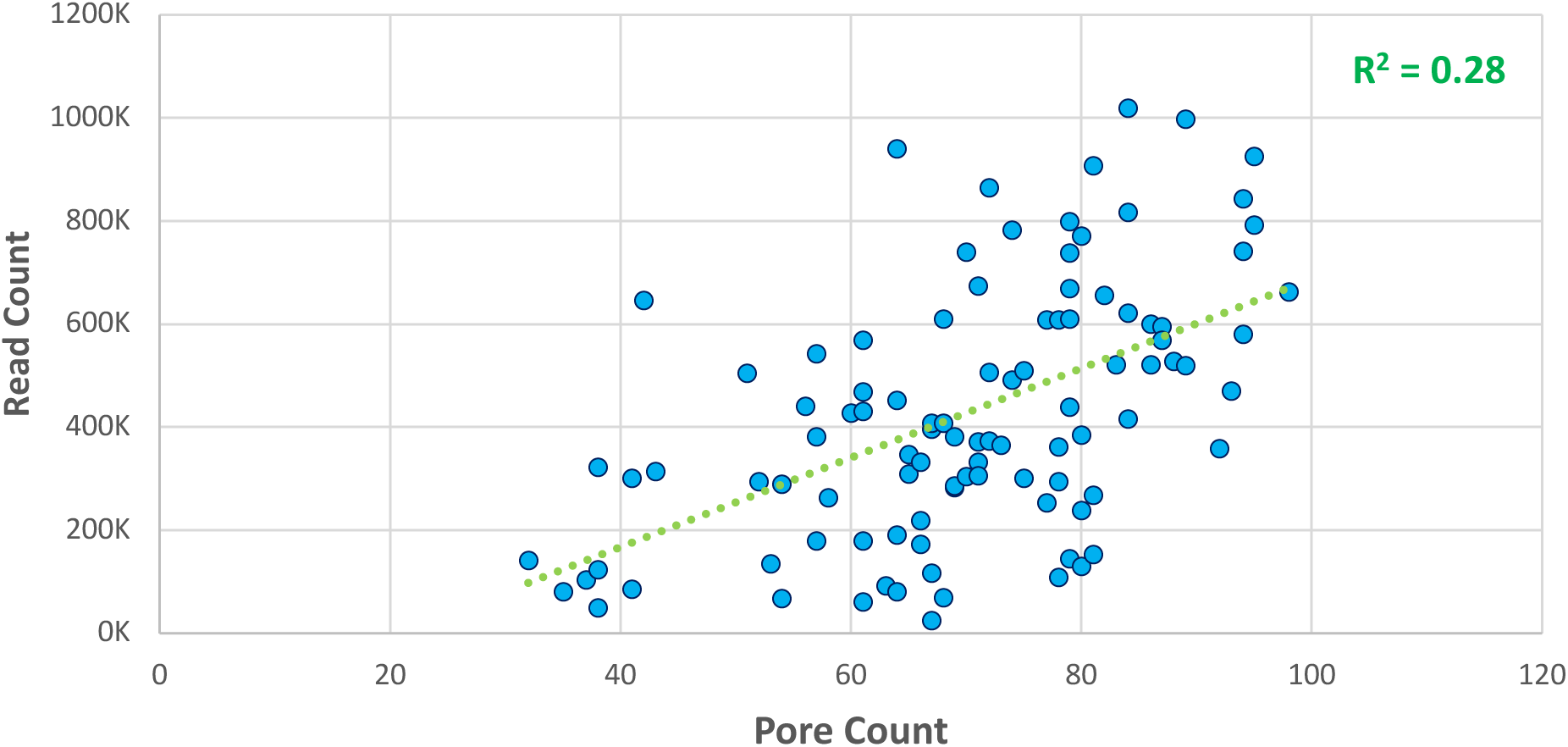
Relationship between the pore count of a Flongle flow cell and reads generated after 24 hours

**Figure 3:**
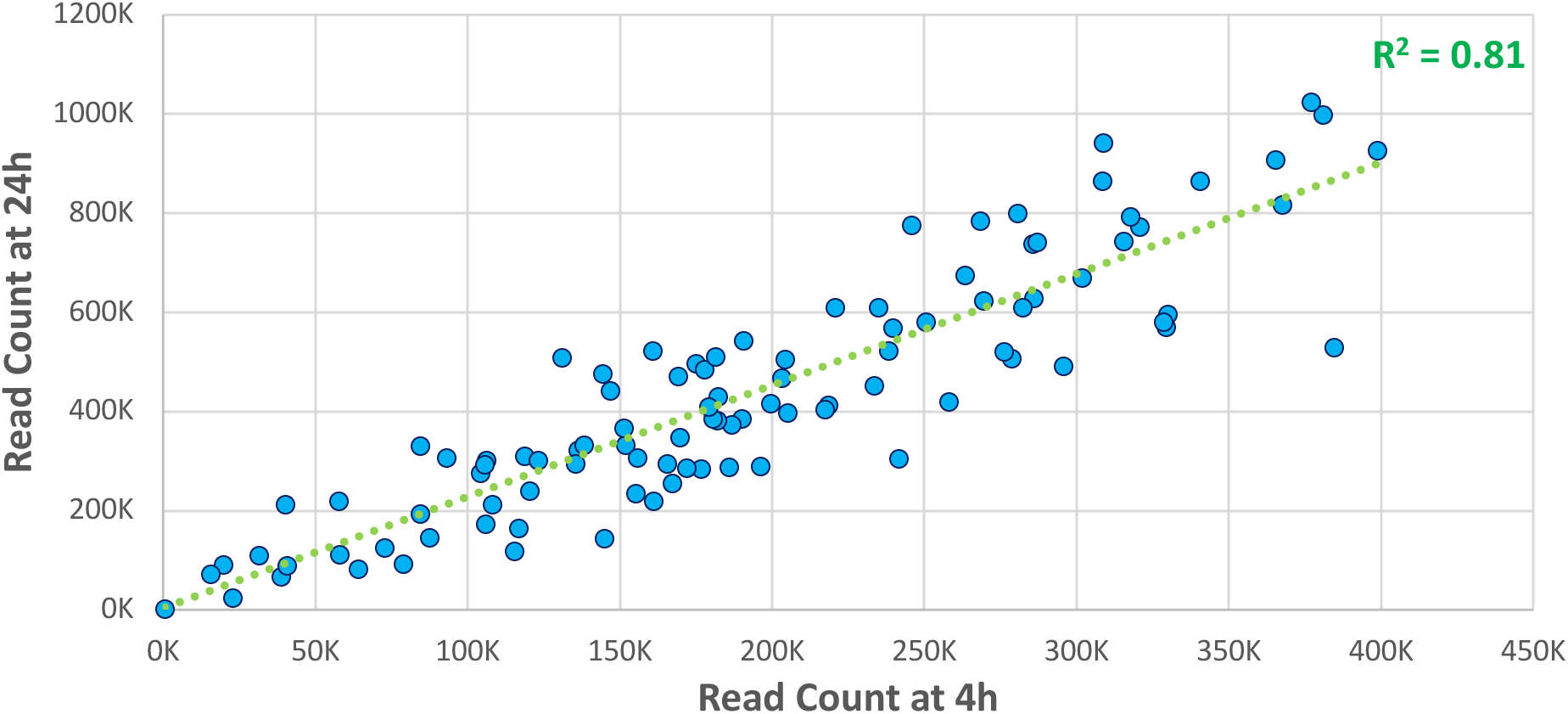
Relationship between read count at 4 and 24 hours based on the analysis of results from 100 Flongle flow cells.

ONT sequences were compared with those generated by SMRT sequencing (99.9% CCS) to ascertain the number of ONT reads required to generate a consensus sequence with less than 0.10% mean divergence from the SMRT reference (**Figure 4**). This analysis showed that consensus sequences based on two ONT reads showed a mean divergence of 1.37% from their SMRT counterpart. However, divergences dropped rapidly, averaging 0.03% when based on five reads. Although this value was well below our target (0.10%), we adopted five reads as the minimum count required to build a consensus. With this information and adopting these criteria, we assessed the capacity of ONT flow cells to recover barcodes from the three pools.

**Figure 4:**
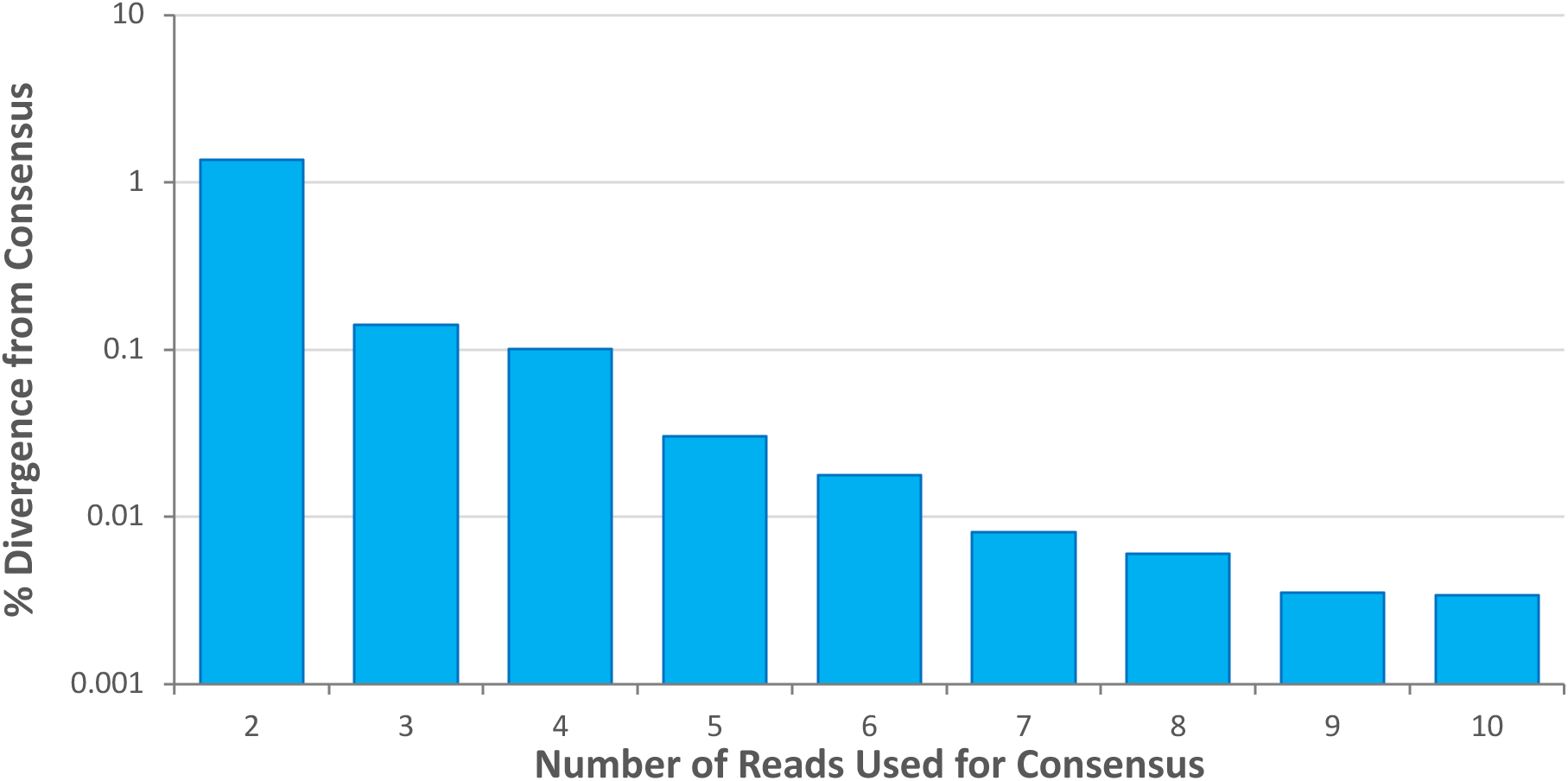
Plot of the log_10_ mean divergence between sequences generated by SMRT analysis on Sequel and sequences based on a variable number of reads from Flongle (10.4.1) flow cells. This analysis examined 1000 specimens from the 10K pool that only generated a single sequence when analyzed on Sequel.

### BARCODE 2K

Analysis of the 2K library on a single Flongle flow cell yielded 939K base-called reads, well above the target (200K). A small loss (6-9%) in read count occurred during size filtering, primer trimming, and assigning reads to the dominant contig (**Figure 5**). By contrast, 42% of the reads failed to meet the criterion for UMI assignment and 36% were not assigned to a contig with 5+ members. As a result, just 283K (30%) of the original reads were retained in the dominant contigs. Inspection of the retained contigs indicated that a barcode from the target (hereafter “on-target sequence”) was recovered from 87% of the specimens (1,987/2,280) with an average of 121 reads each (**Figures 5 & 6A**).

**Figure 5:**
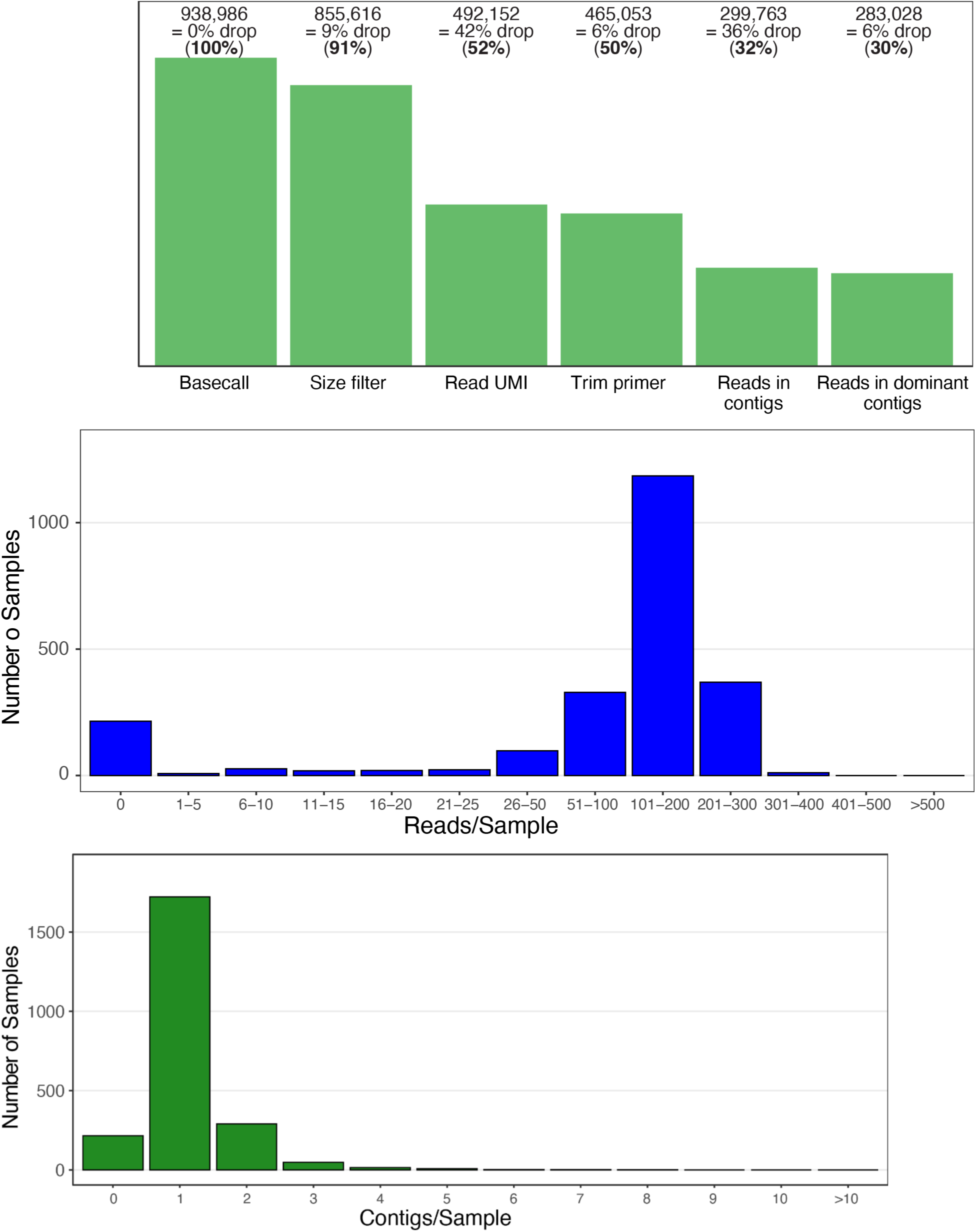
Histograms for the 2K pool showing the read loss at each step in the bioinformatic pipeline (remaining proportion is shown in parentheses), the number of reads per sample, and the number of contigs per sample.

Among all specimens, 72% delivered only an on-target sequence, 15% delivered both on- and off-target sequences, while 9% delivered no sequence, and 4% only had an off-target sequence (**Supplemental Table 2**). Most off-target sequences were either endosymbiotic bacteria (Rickettsiales) or were unassigned because they had >40% divergence to any COI sequence on BOLD. In cases where they occurred together with an on-target sequence, they generally had a lower read count. Although most specimens only delivered a single contig, 16% delivered 2 or 3 contigs and the others produced up to 8 contigs (**Figure 5**).

Considering all insects (n = 2,031), sequence recovery averaged 87%, but varied by 24% among the orders (93%–Lepidoptera, 93%–Hemiptera, 91%–five small orders, 90%–Diptera, 81%–Hymenoptera, 69%– Coleoptera) (**Table 2**). Members of the other two classes (Arachnida = 134; Collembola = 115) had success rates of 77% and 91% respectively. Sequence recovery was correlated with mean read depth per specimen for on-target sequences (**Supplemental Figure 1**) which was 123 for insects, 99 for arachnids, and 110 for collembolans (**Table 2**).

**Table 2:** Number of specimens from 28 orders of arthropods in the 2K, 10K, and 100K pools and percent success in sequence recovery from them. A few specimens could not be assigned to class (12) or order (14) because they lacked an image, but most were likely Mesostigmata.

| Class | Order | 2K |  | 10K |  | 100K |  |
| --- | --- | --- | --- | --- | --- | --- | --- |
|  |  | n | Success | n | Success | n | Success |
| unknown | unknown | - | - | - | - | 12 | 17% |
| Arachnida | unknown | 9 | 22% | 9 | 0% | 14 | 0% |
| Arachnida | Araneae | 33 | 76% | 33 | 67% | 134 | 76% |
| Arachnida | Mesostigmata | 17 | 71% | 17 | 59% | 22 | 73% |
| Arachnida | Opiliones | - | - | - | - | 3 | 67% |
| Arachnida | Pseudoscorpiones | - | - | - | - | 3 | 0% |
| Arachnida | Trombidiformes | 75 | 88% | 76 | 82% | 84 | 92% |
| Collembola | Entomobryomorpha | 112 | 91% | 112 | 94% | 116 | 97% |
| Collembola | Symphyleona | 3 | 100% | 3 | 33% | 5 | 60% |
| Insecta | Archaeognatha | - | - | - | - | 8 | 50% |
| Insecta | Blattodea | - | - | - | - | 144 | 97% |
| Insecta | Coleoptera | 119 | 69% | 307 | 70% | 4533 | 81% |
| Insecta | Dermaptera | - | - | - | - | 7 | 86% |
| Insecta | Diptera | 696 | 90% | 4603 | 92% | 76084 | 93% |
| Insecta | Ephemeroptera | - | - | - | - | 48 | 8% |
| Insecta | Hemiptera | 243 | 93% | 489 | 83% | 3854 | 83% |
| Insecta | Hymenoptera | 480 | 81% | 1726 | 78% | 9418 | 84% |
| Insecta | Lepidoptera | 352 | 93% | 1452 | 96% | 4448 | 95% |
| Insecta | Mantodea | - | - | - | - | 11 | 73% |
| Insecta | Neuroptera | 41 | 88% | 83 | 83% | 103 | 82% |
| Insecta | Odonata | - | - | - | - | 2 | 0% |
| Insecta | Orthoptera | 1 | 0% | 1 | 0% | 165 | 81% |
| Insecta | Parasitidae | 1 | 100% | 1 | 0% | 1 | 100% |
| Insecta | Phasmatodea | - | - | - | - | 2 | 50% |
| Insecta | Psocodea | 6 | 67% | 19 | 58% | 443 | 39% |
| Insecta | Siphonaptera | - | - | 1 | 100% | 1 | 100% |
| Insecta | Strepsiptera | - | - | - | - | 1 | 100% |
| Insecta | Thysanoptera | 89 | 94% | 177 | 94% | 209 | 93% |
| Insecta | Trichoptera | 3 | 67% | 11 | 55% | 444 | 78% |
| Insecta | Zoraptera | - | - | - | - | 1 | 0% |

All 1,987 on-target barcode sequences fell within the expected length range (643–661 bp) and 93% had no ambiguous base calls (**Supplemental Table 3**; **Supplemental Figure 2**). Considering both platforms, sequences were recovered from 93.3% of the specimens (2,127/2,280) with 3% higher success on Sequel (2,049) than ONT (1,987) (**Figure 6A**). However, both platforms recovered sequences from many of the same specimens (89.8%). A fidelity check indicated that 94% of the ONT sequences were identical to their SMRT counterpart while 98% showed less than 2% divergence (**Figure 6B**).

**Figure 6:**
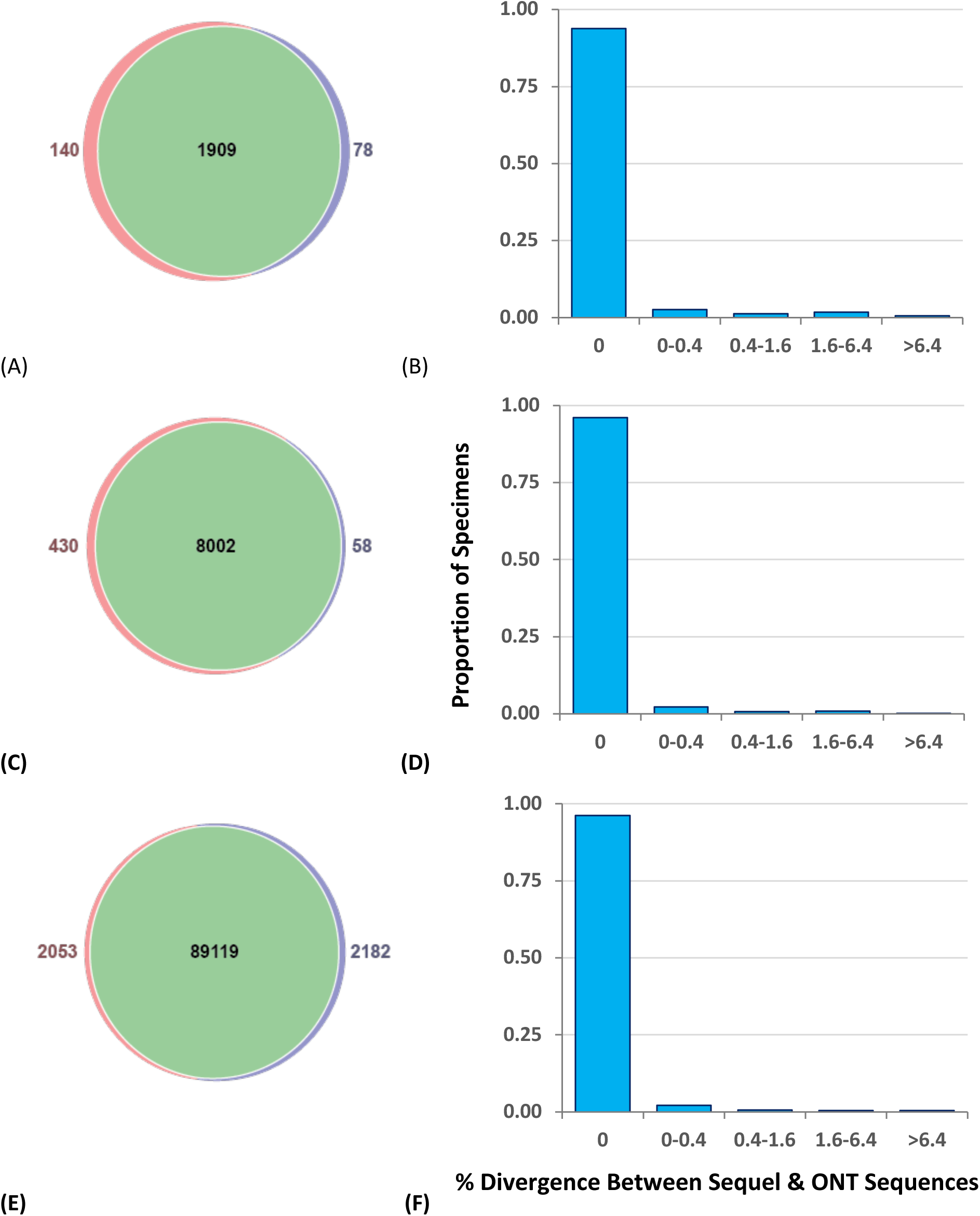
Venn diagrams showing the number of specimens from the 2K (A), 10K (B), and 100K (C) libraries that yielded a barcode sequence on both platforms (green), only on Sequel (pink), or only on ONT (blue). Histograms showing the proportion of specimens yielding sequences on both ONT and Sequel in three pools (B = 2K, D = 10K, F = 100K) that fell into five divergence categories.

### BARCODE 10K

As the 1M target was unlikely to be achieved with a single Flongle flow cell, read count was assessed after four hours, and this library was loaded onto two more flow cells, a strategy which yielded 1.2M base-called reads. A small loss (6-11%) in read count occurred during size filtering, primer trimming, and assigning reads to the dominant contig (**Figure 7**). By contrast, 46% of the reads failed to meet the criterion for UMI assignment and 41% were not assigned to a contig with 5+ members. As a result, just 331K (26%) of the original reads were retained in the dominant contigs. Inspection of these retained contigs indicated that on-target sequences were recovered from 88.4% of the specimens (8,060/9,120) with a mean of 40 reads (**Figures 6C & 7**).

**Figure 7:**
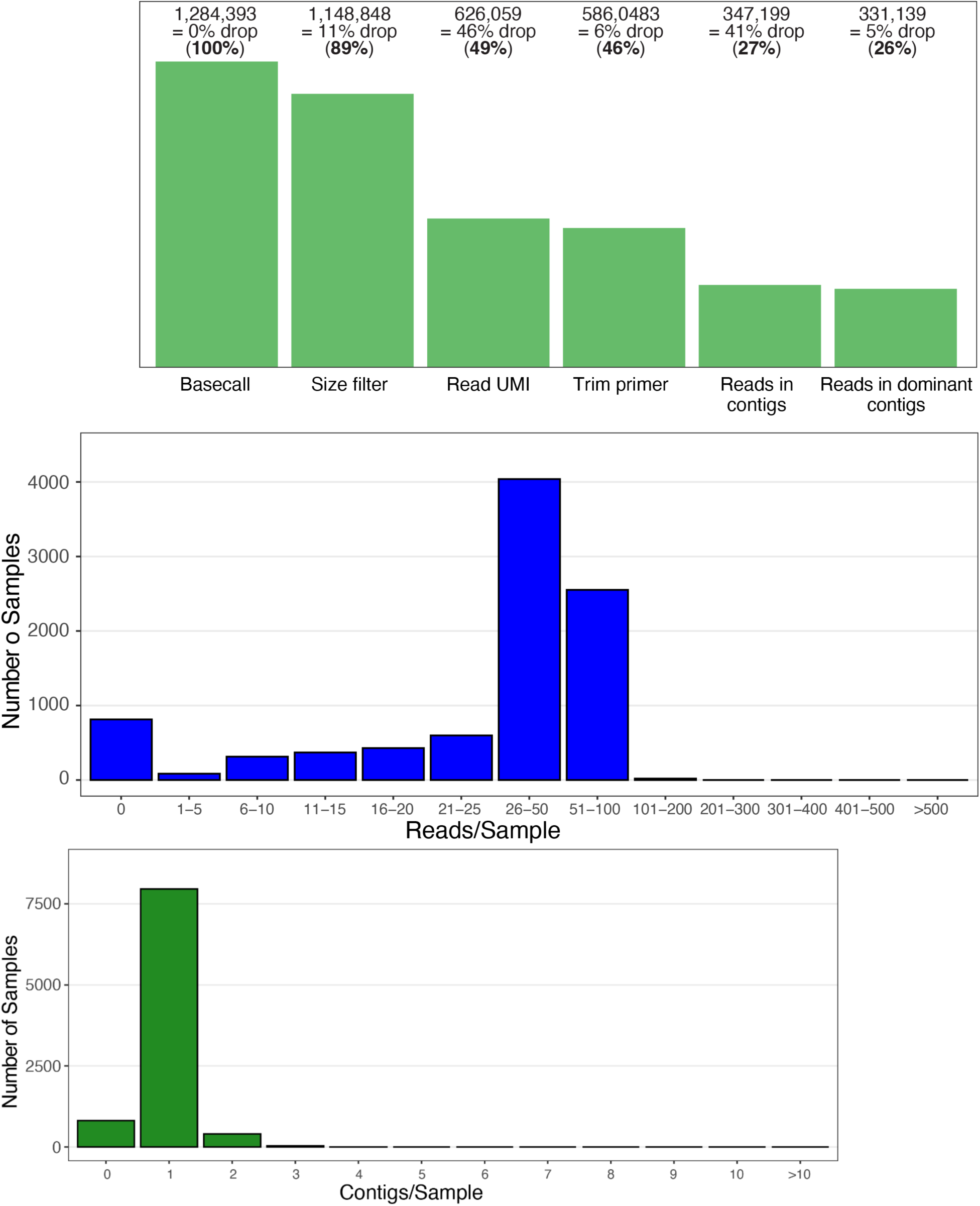
Histograms showing the read loss at each step in the bioinformatic pipeline (remaining proportion is shown in parentheses) for the 10K library, the number of reads per sample, and the number of contigs per sample.

Among all specimens, 84% delivered only an on-target sequence, 5% both on- and off-target sequences, 8% no sequence, and 3% only an off-target sequence (**Supplemental Table 2**). Although most specimens only delivered a single contig, 5% delivered 2 or 3 contigs and the others produced up to 4 contigs (**Figure 7**).

Barcodes were recovered from 90% of the insects (n = 8,869) with recovery varying by 17% among the orders (96%–Lepidoptera, 93%–Diptera, 93%–Coleoptera, 87%–six small orders, 82%–Hemiptera, 79%– Hymenoptera) (**Table 2**). Arachnida (n = 135) and Collembola (n = 115) had success rates of 72% and 92% respectively. Sequence recovery was correlated with mean read depth per specimen for on-target sequences (**Supplemental Figure 1**) with 36 for insects, 24 for arachnids, and 31 for collembolans (**Table 2**). 99.9% of the 8,060 on-target barcodes fell within the expected length range (643–661 bp), and 94% lacked any ambiguous base calls (**Supplemental Table 4**; **Supplemental Figure 2**). Considering both platforms, sequences were recovered from 93.1% of the specimens (8,490/9,120) with 4.6% higher success on Sequel (8,432) than ONT (8,060) (**Figure 6C**). However, both platforms recovered sequences from many of the same specimens (94.3%). A fidelity check indicated that 96% of the ONT barcodes were identical to their SMRT counterpart while 99% showed less than 2% divergence (**Figure 6D**).

### BARCODE 100K

The MinION flow cell generated 19M base-called reads for the 100K library, comfortably exceeding the target (10M). These sequences went through a 6-step filtration process because of the additional level of demultiplexing linked to the use of both UMIs and Native barcodes. A small loss (4-9%) in read count occurred during native barcode assignment, size filtering, primer trimming, and assigning reads to the dominant contig (**Figure 8**). More read loss was observed at the other stages; 30% failed to meet the criterion for UMI assignment and 14% were not assigned to a contig with 5+ members. Despite these losses, 9.1M (47%) of the original reads were retained in the dominant contigs (**Figure 8**). Inspection of the retained contigs indicated that sequences were recovered from 92% of the specimens (91,301/100,320) with an average of 97 reads (**Figures 6E & 8**) with 80% delivering an on-target sequence, 12% both on- and off-target sequences, 6% no sequence, and 2% only an off-target sequence (**Supplemental Table 2**). Although most specimens only delivered a single contig, 13% delivered 2 or 3 contigs and the others produced up to 8 contigs.

**Figure 8:**
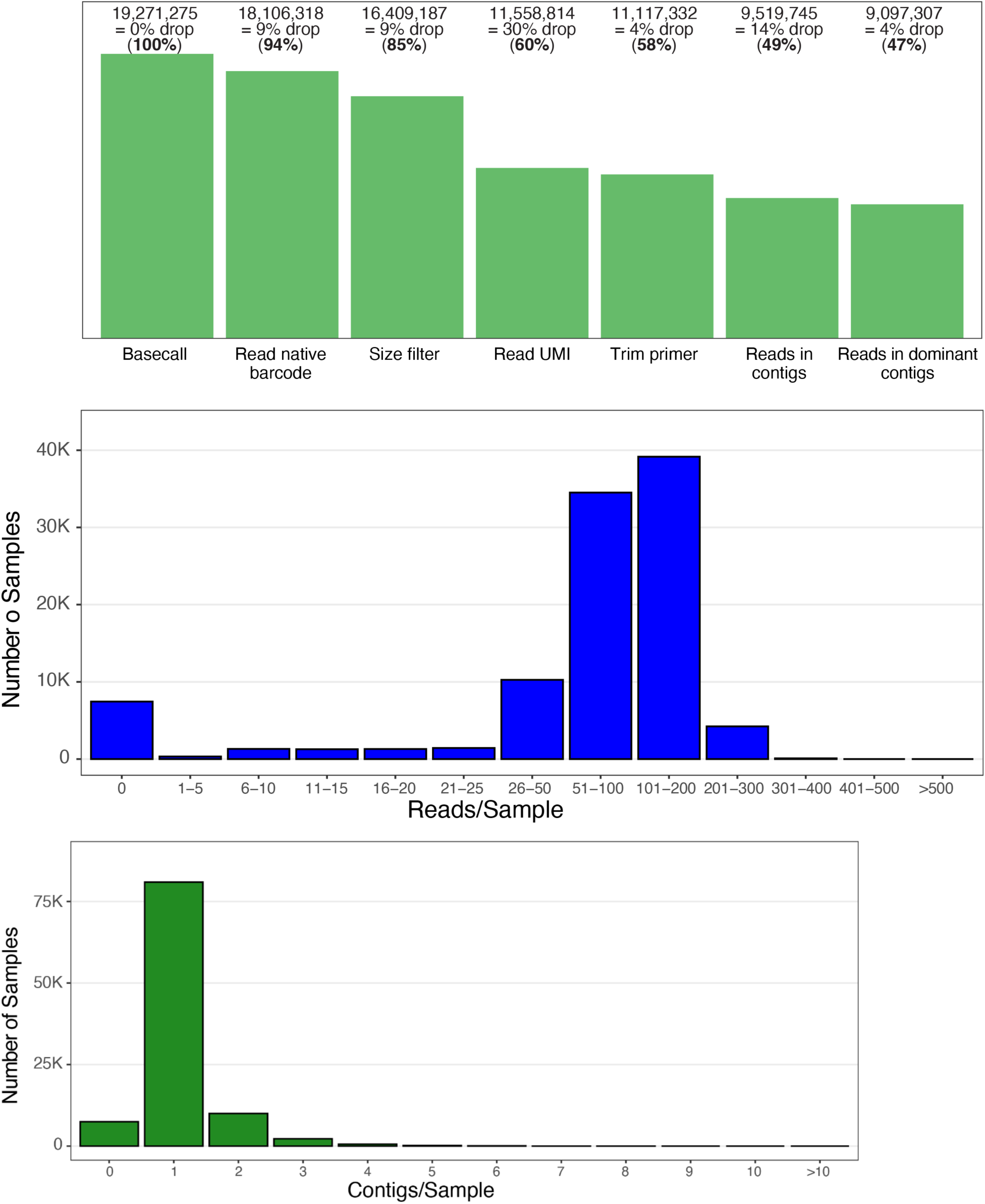
Histograms showing the read loss at each step in the bioinformatic pipeline (remaining proportion is shown in parentheses) for the 100K library, the number of reads per sample, and the number of contigs per sample.

The 99,920 insects had an overall success rate of 92% with recovery varying by 25% among the orders (95%–Lepidoptera, 94%–Diptera, 85%–Hymenoptera, 84%–Hemiptera, 84%–Coleoptera, 70%–15 small orders) (**Table 2**). Arachnida (n = 260) and Collembola (n = 121) had overall success rates of 78% and 95% respectively. Sequence recovery was correlated with mean read counts/specimen for on-target sequences (**Supplemental Figure 1**) which was 90 for insects, 66 for arachnids, and 106 for collembolans (**Table 2**).

99.5% of the 91,301 on-target ONT sequences fell within the expected length range (643–661 bp) and 97% lacked any ambiguous base calls (**Supplemental Table 5**; **Supplemental Figure 2**). Considering both platforms, sequences were recovered from 92.1% of the specimens (93,354/101,320) with 0.1% higher success on ONT (91,301) than Sequel (91,172) (**Figure 6E**). However, both platforms recovered sequences from many of the same specimens (95.4%). A fidelity check indicated that 96% of the ONT barcodes were identical to their SMRT counterpart while 99% showed less than 2% divergence (**Figure 6F**).

## DISCUSSION

This study has assessed the capacity of the latest generation of ONT flow cells (R10.4.1) to generate high-fidelity DNA barcode records in a cost-effective and scalable fashion. As such, we tested sequence recovery across three levels of amplicon pooling (2K, 10K, 100K specimens). Before examining recovery success, we set a fidelity criterion, one requiring ONT sequences to have a mean sequence divergence that was less than 0.1% from sequences generated on Sequel/Sequel II using a 99.9% CCS. Our analysis indicated that five ONT reads were required to meet this criterion so any specimen yielding fewer sequences was excluded from recovery, even if it possessed 3–4 identical sequences. The adoption of this stringent criterion reduced sequence recovery but ensured those retained were effectively identical to their Sequel counterparts. Aside from this fidelity criterion, we set a target of 100 reads per specimen to aid sequence recovery in the face of the inescapable variation in product concentration across the PCR reactions combined in each pool and the random sampling of amplicons during sequencing.

Having set a quality standard and target read depth, we focused on identifying other factors that might complicate the use of ONT for DNA barcoding. We directed these efforts towards the Flongle flow cell as its R10.4.1 version is in beta while MinION has completed validation. Our investigation revealed no evidence of a decline in flow cell performance over five months, 5x the warrantied expiration (one month), but a tenth of the Flongles failed to meet the warrantied pore count. Among those achieving this standard, we observed 40-fold variation in the number of reads generated. As expected, this variation was positively associated with pore count, but the correlation was weak. Examination of the number of reads generated by each pore explained the lack of a strong correlation as pores varied tremendously in sequence output. For example, the 895 pores in a MinION flow cell generated from 1 to 22,000 reads (**Supplemental Figure 3**). The variation in read counts generated by different Flongle flow cells introduced little complexity in meeting our target read depth. Because the read count after four hours was a good predictor of the final count, the production of each Flongle was tracked, and sequencing depth was adjusted by loading additional flow cell(s) as required. With this approach, we reliably met our target of 100 base-called reads per specimen. The deployment of additional flow cells was straightforward because the standard ONT library protocol typically generates enough template to load several Flongles.

In our study, Flongle flow cells produced a mean of 0.45M base-called reads whereas MinION generated roughly 40x more, reflecting its higher pore count and run time. 30% of the reads generated by Flongles were assigned to a dominant contig, while 47% of those produced by the MinION met this criterion despite the fact that data analysis involved an extra step. Both flow cells showed low loss of sequences during primer trimming, size filtering, and assignment to dominant contigs. The major loss steps occurred during UMI categorization, a step that required no more than two nucleotide changes at either end of the amplicon and in assigning reads to a contig. Low sequence fidelity would reduce success in UMI assignment and would create more low frequency sequences that would not be assigned to a contig. Although more study is required, the R10.4.1 version of MinION appeared to deliver higher fidelity sequences than its Flongle counterpart because UMI assignments experienced 50% less loss with MinION, and 50% more of the sequences were assigned to a dominant contig.

Earlier studies have demonstrated that ONT can support barcode analysis, but they involved pooling amplicons from < 250 specimens (Cuber et al. 2023, Srivathsan et al. 2021). Our study breaks ground by showing that a single Flongle flow cell can support the analysis of 2K specimens, while two or three combined can characterize 10K, and a MinION flow cell can analyze 100K. More than 95% of the sequences recovered in our studies showed identity with high-fidelity sequences generated on Sequel/Sequel II. Although >98% of ONT sequences were 658 bp, others ranged from 643–661 bp, reflecting codon insertions/deletions in the barcode region of some insect lineages (Pentinsaari et al. 2016).

Sequel II can generate barcode sequences for $0.05 per specimen, but this requires the analysis of 50K pools (pers. obs.). By comparison, ONT can deliver high-fidelity sequences at $0.05 per specimen for a 2K pool, declining to $0.01/specimen for a 100K pool. A further reduction should be achievable as amplicons from 884,736 specimens can be separately tagged by coupling 96F x 96R UMIs with ONT’s 96 native barcodes. A PromethION flow cell would deliver the desired target read count (100M) while reducing sequencing costs to $0.001/specimen. Because ONT avoids the high capital and service costs of SequeI platforms and delivers sequences inexpensively across diverse pool sizes, it is well suited to support the activation of a global network of barcode centres. Moreover, the low cost of its base flow cells will be a stimulant for protocol development.

Aside from delivering sequences inexpensively, ONT closely matched the capacity of Sequel to recover barcodes from each pool. Prior extensive work on Sequel/Sequel II has shown that it is difficult to recover sequences from more than 90% of the specimens in any highly multiplexed pool of arthropods (Hebert *et al*. 2018). In part, this reflects the differential amplicon abundances in the pool and the random sampling of these templates during sequencing. However, 100% recovery is additionally impeded because standard primers fail to bind to the barcode region of some species. This problem is most acute in hymenopteran lineages with accelerated rates of mitochondrial evolution (Kaltenpoth *et* al. 2012), a fact reflected in lower barcode recovery for this order in this and in earlier studies (Hebert et al. 2016).

Reflecting a 20-year effort, BOLD now holds 15 million DNA barcode records derived from 1.2 million species of eukaryotes (Ratnasingham and Hebert 2007). Over this interval, production rates have risen with iBOL’s largest analytical facility now processing 3 million specimens annually. In looking to the future, the effort required to gain DNA barcode coverage for all species is uncertain because the species count is unknown. Mora et al. (2012) estimated the global species count at 8.7M species, but this is certainly too low, reflecting serious underestimates of species richness in some hyper-diverse lineages. Although arthropods remain the most diverse phylum of multicellular life, Diptera and Hymenoptera (Hebert et al. 2017, Forbes et al. 2018) have dislodged Coleoptera from its long-standing position as the most diverse insect order (Stork et al. 2015; Stork 2018). This reevaluation, coupled with new evidence that other arthropod lineages such as mites (Young et al. 2019) include millions of species, reinvigorates earlier suggestions that there could be 30 million species of arthropods (Erwin 1982). As the Nematoda (Hodda 2022) and fungi (Baldrian et al. 2022) likely each include another 10 million species, Wiens (2023) suggested there could be 100 million species of eukaryotes.

Because most species are undescribed, comprehensive DNA barcode coverage cannot be gained by targeting known species lacking representation on BOLD. Instead, the goal of comprehensive coverage can only be achieved by sequencing massive numbers of specimens and using the BIN system to register the species encountered. This brute force approach to biodiversity characterization means that the number of analyses will necessarily far exceed the species count. BOLD currently holds 12x more records than species and this ratio will certainly rise as work advances, meaning that completion of a barcode reference library for all eukaryotes will likely require analyzing more than a billion specimens. Given this count, it is critical to identify inexpensive analytical protocols. As ONT has effectively solved the challenge for sequencing, similar economies for DNA extraction and PCR must be sought.

## Supporting information

Supplemental Table 1 - Specimen Details

Supplemental Table 2 - Contig and read summary

Supplemental Table 3 - Dominant Contigs 2K

Supplemental Table 4 - Dominant Contigs 10K

Supplemental Table 5 - Dominant Contigs 100K

## ACKNOWLEDGEMENTS

This work was funded by the Government of Canada through Genome Canada and Ontario Genomics (OGI-208, OGI-233), by the New Frontiers in Research Fund (NFRFT-2020-00073), and by the Canada Foundation for Innovation.

---

**Supplemental Figure 1:**
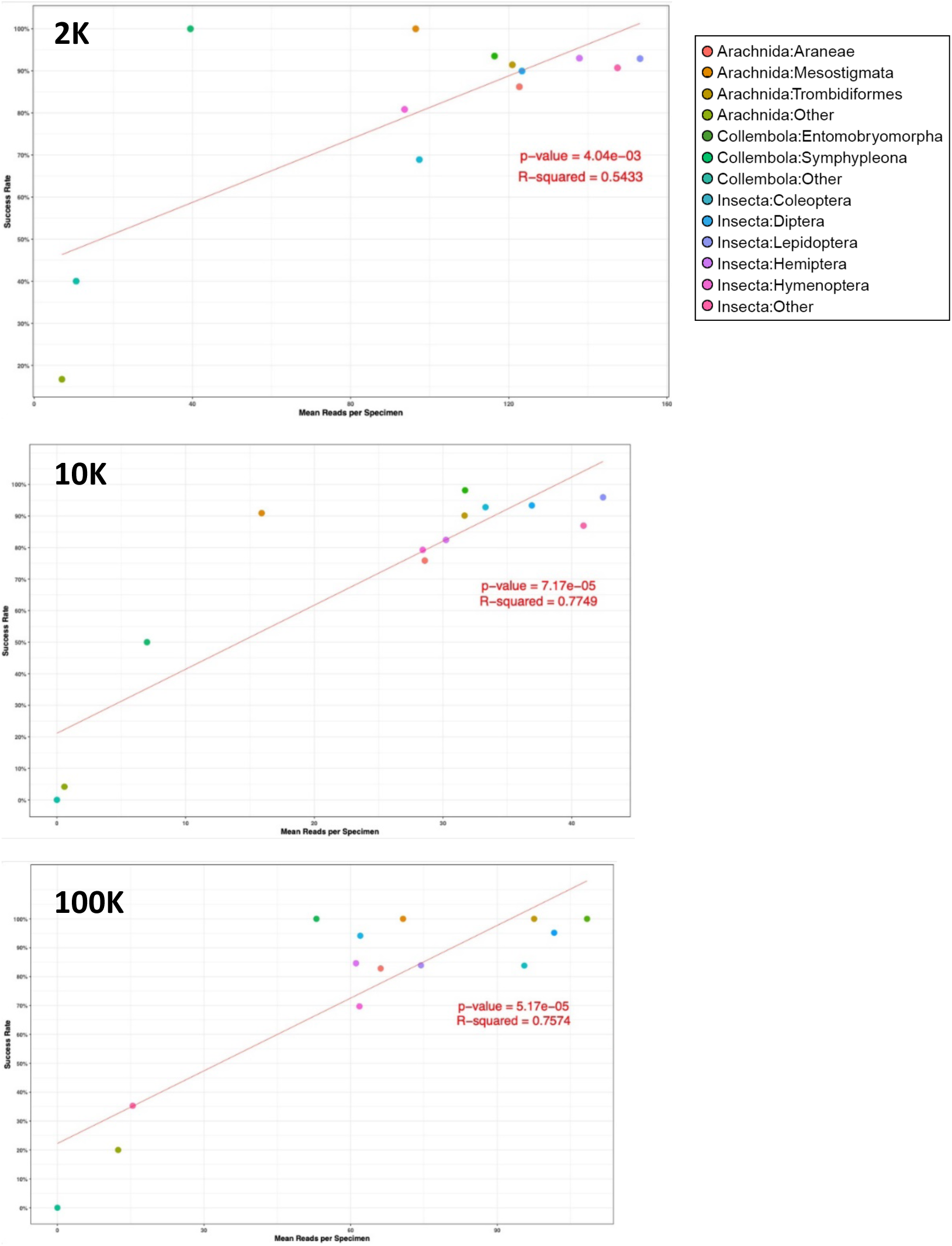
Correlation between the mean number of reads per specimen and sequence recovery for 13 arthropod groups in the 2K, 10K, and 100K libraries.

**Supplemental Figure 2:**
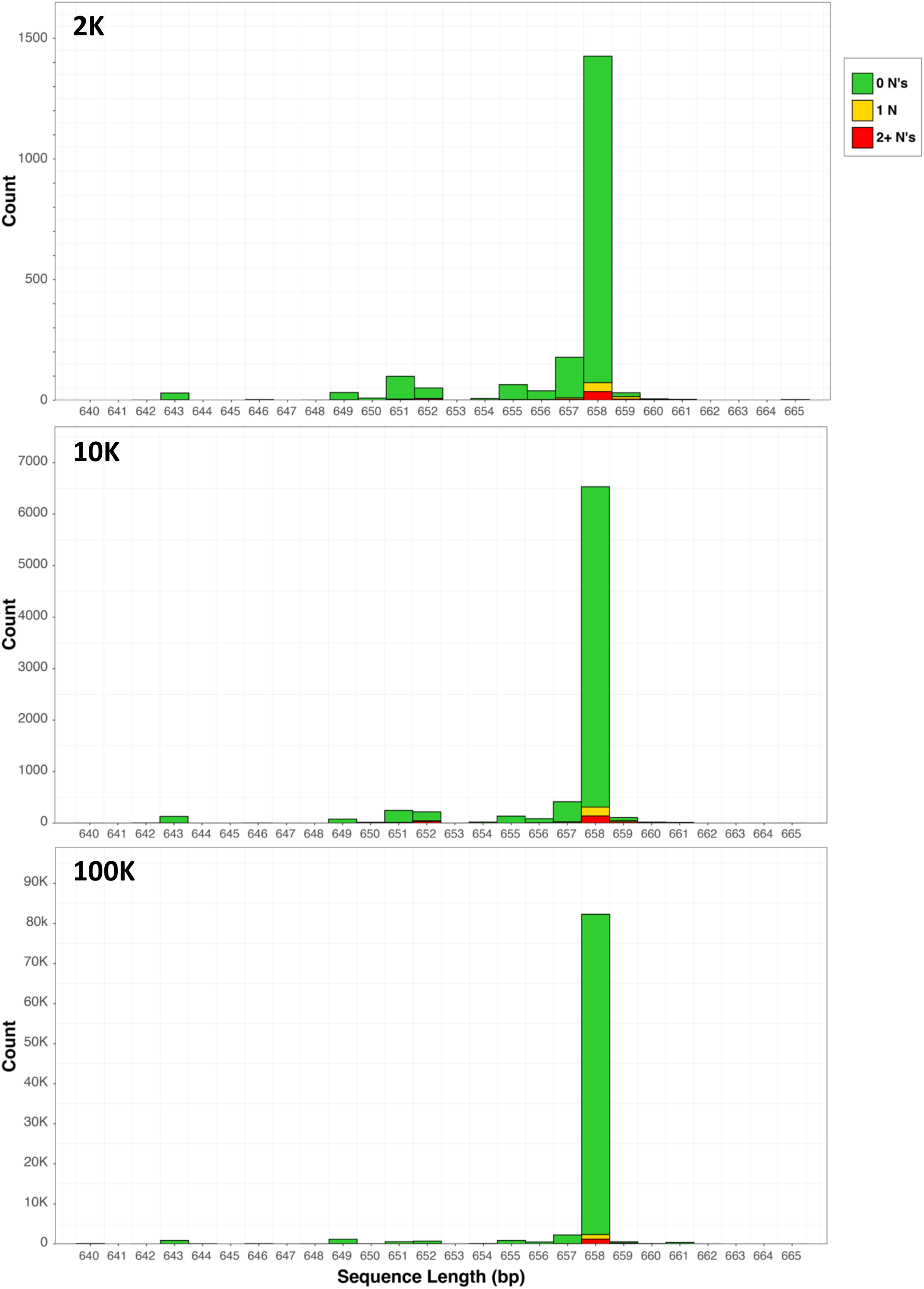
Histograms showing variation in lengths of ONT barcodes recovered from the 2K, 10K, and 100K pools. Colours discriminate records with 0, 1, or 2+ ambiguous base calls.

**Supplemental Figure 3:**
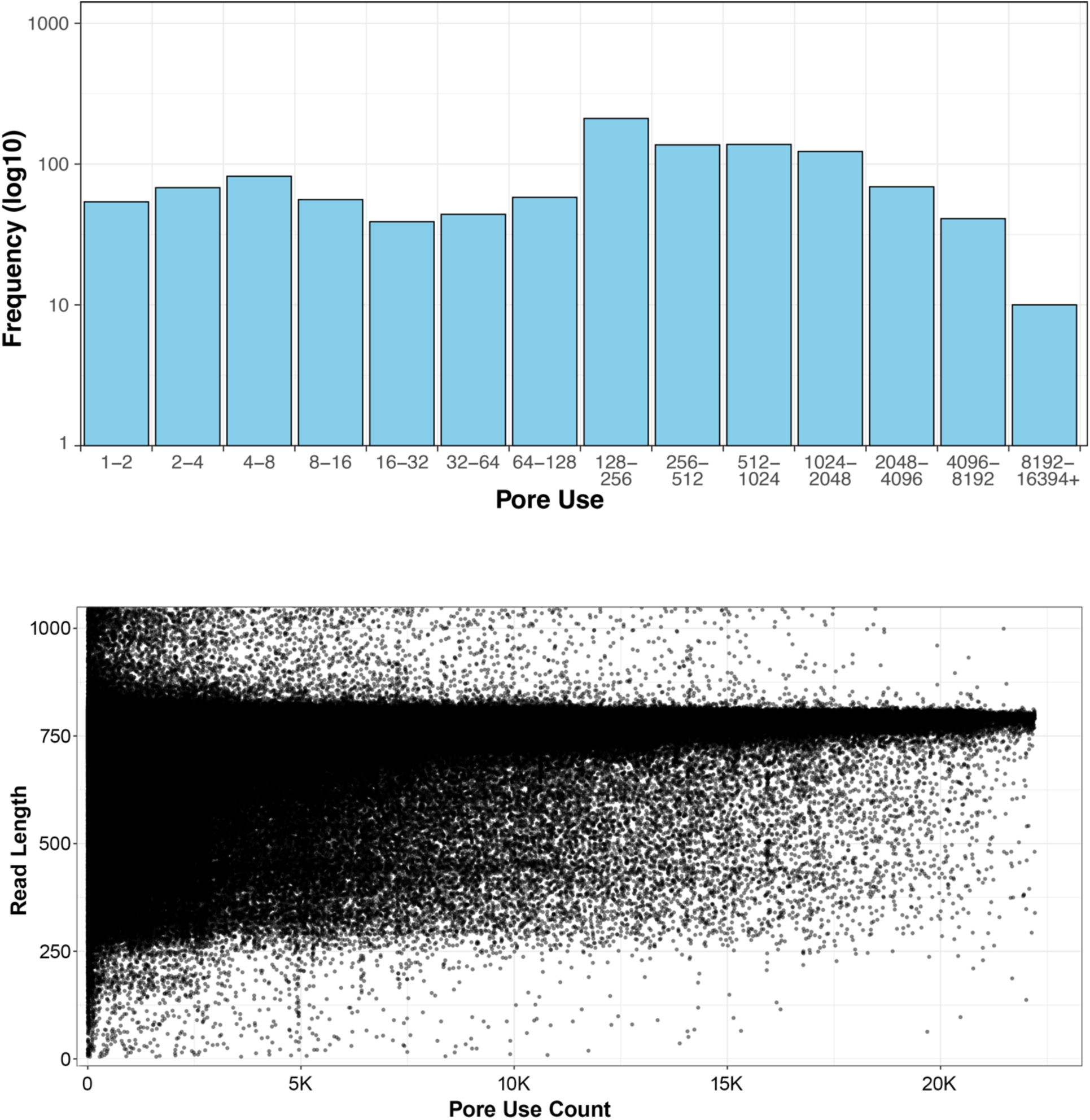
Histogram showing the number of sequences generated by each of the 895 active pores in a MinION flow cell. Scatterplot showing the length of a subset of 1M reads generated by this flow cell. Sequences vary from 1-1,100 bp, but cluster near the length of the target COI amplicon together with UMIs and primers (770 bp). The x-axis shows when a particular sequence was generated in relation to others in the same run.

**Supplemental Table 1:** *BOLD Sample IDs and taxonomic details of all 100,300 specimens included in this study. Whether or not each of the specimens was included in the 2K, 10K, or 100K experiments is also stated.*

**Supplemental Table 2:** *Contig and read counts for every specimen in each of the three experiments (2K, 10K, 100K).*

**Supplemental Table 3:** *Details of each on-target sequence yielded in the 2K experiment. For each sequence, the nucleotide sequence itself is provided, along with sequence length, read depth, the number of ambiguous bases, and the source specimen Sample ID and taxonomy.*

**Supplemental Table 4:** *Details of each on-target sequence yielded in the 10K experiment. For each sequence, the nucleotide sequence itself is provided, along with sequence length, read depth, the number of ambiguous bases, and the source specimen Sample ID and taxonomy.*

**Supplemental Table 5:** *Details of each on-target sequence yielded in the 2K experiment. For each sequence, the nucleotide sequence itself is provided, along with sequence length, read depth, the number of ambiguous bases, and the source specimen Sample ID and taxonomy.*

