## Supplemental File 1 for "Barcode 100K Specimens: In a Single Nanopore Run": autotrim_taxfilter_dominants.R

```
#get argument from bash script (plate name)
args <- commandArgs(trailingOnly = TRUE)
plateid <- args[1]
BOLDtax <- args[2]

#set working directory based on plate's folder, load required libraries
setwd(paste("/home/innovation-admin/DATA_INPUT/",plateid,"/", sep = ""))
library("Biostrings")
library("readxl")
library(muscle)

#input contig consensus sequences and tax ID table
input.fasta <- readDNAStringSet(sprintf("%s.fasta", plateid))
input.table <- read.table(sprintf("%s.table", plateid), header = TRUE, fill = TRUE, row.names = NULL, sep = "\t")
input.table <- input.table[-1,-5]
names(input.table) <- c("SeqName","TaxAssign","Strand","TaxAssignFinal")

#### auto-trim contig consensus sequences ##### *** INPUT FASTA MUST HAVE AT LEAST 100 SEQUENCES ***
num_seqs <- length(input.fasta)
num_full_batches <- floor(num_seqs / 100)
num_last_batch <- num_seqs %% 100
if (num_last_batch != 0){
  last_batch <- num_full_batches + 1
} else {
  last_batch <- num_full_batches
}
second_last_batch <- last_batch - 1
batches <- list()

#if fewer than 100 sequences, skip auto-trimming
if (num_seqs < 100){
  output.fasta <- input.fasta
} else {
  #split FASTA file into batches of 100 sequences + last remaining sequences
  for (i in 1:last_batch) {
    if(i == second_last_batch){
      start <- (i - 1) * 100 + 1
      end <- num_seqs
      batch <- input.fasta[start:end]
      batches <- c(batches, list(batch))
    } 
    if(i != second_last_batch & i != last_batch) {
      start <- (i - 1) * 100 + 1
      end <- i * 100
      batch <- input.fasta[start:end]
      batches <- c(batches, list(batch))
    }
  }

  #align each batch and trim ends
  output.fasta <- DNAStringSet()
  for (q in batches) {
    #align sequences
    aligned <- DNAStringSet(muscle(q))
    
    #get start and end positions
    column_counts <- colSums(as.matrix(aligned) == "-")
    columns_with_fewer_than_95p_gaps <- which(column_counts < 0.05*length(aligned))
    first_col <- as.numeric(columns_with_fewer_than_95p_gaps[1])
    last_col <- as.numeric(columns_with_fewer_than_95p_gaps[length(columns_with_fewer_than_95p_gaps)])
    
    #trim alignment to start and end positions, then un-align and add to master FASTA file
    trimmed_alignment <- subseq(aligned, start = first_col, end = last_col)
    unaligned <- DNAStringSet(sapply(trimmed_alignment, gsub, pattern = "-", replacement = "", USE.NAMES = FALSE))
    output.fasta <- c(output.fasta, unaligned)
  }
}
writeXStringSet(output.fasta,sprintf("%s_AllContigs.fasta", plateid))
input.fasta <- output.fasta

##### add sequences (auto-trimmed if performed) to contig tax id table #####
names <- names(input.fasta)
seqs <- as.character(input.fasta, use.names = FALSE)
df <- data.frame(Sequence = seqs, SeqName = names, stringsAsFactors = FALSE)
parsed <- strsplit(df$SeqName, "\\|")
df$Sample <- sapply(parsed, "[[", 1)
df$ContigName <- sapply(parsed, "[[", 2)
df$ReadCount <- sapply(parsed, "[[", 3)
df <- df[,c(3,4,5,1,2)]
df$ReadCount <- as.numeric(as.character(gsub("reads-","",df$ReadCount)))
df <- df[order(df$Sample, -df$ReadCount),]
output <- merge(df, input.table, by = "SeqName", all.x = TRUE)
split_values <- strsplit(output$TaxAssignFinal, ",")
#max_length <- max(lengths(split_values))
max_length <- 6
matrix_values <- t(sapply(split_values, function(x) c(x, rep(NA, max_length - length(x)))))
new_cols <- as.data.frame(matrix_values)
output <- cbind(output, new_cols)
output[is.na(output)] <- "unknown"
names(output)[9:14] <- c("Phylum","Class","Order","Family","Genus","Species")
output <- output[,c(2,3,4,9:14,1,5,6,8)]
output$Phylum <- gsub("p:", "", output$Phylum, fixed = TRUE)
output$Class <- gsub("c:", "", output$Class, fixed = TRUE)
output$Order <- gsub("o:", "", output$Order, fixed = TRUE)
output$Family <- gsub("f:", "", output$Family, fixed = TRUE)
output$Genus <- gsub("g:", "", output$Genus, fixed = TRUE)
output$Species <- gsub("s:", "", output$Species, fixed = TRUE)
output$PlateOrRunID <- plateid
output <- output[,c(14,1:13)]

#add sequence length to output table
output$SeqLength <- nchar(output$Sequence)
output <- output[,c(1:4,15,5:14)]

#if BOLD taxonomy is not available, output table as is
if (BOLDtax == "no") {
  #output table
  write.table(output, sprintf("%s_TaxonomicAssignments_AllContigs.tsv", plateid), append = F, quote = F, row.names = F, sep = "\t")
}

#if BOLD taxonomy is available, flag order-level mismatches in contig table then output table
if (BOLDtax == "yes") {
  #generate BOLD tax table based on PIDs and SIDs (in case either was used)
  input.bold.lab.sheet <- data.frame(read_excel(list.files("/home/innovation-admin/DATA_INPUT", pattern = "taxonomy.*\\.xlsx", full.names = TRUE),sheet = "Lab Sheet", skip = 2))
  input.bold.taxonomy <- data.frame(read_excel(list.files("/home/innovation-admin/DATA_INPUT", pattern = "taxonomy.*\\.xlsx", full.names = TRUE),sheet = "Taxonomy", skip = 2))
  input.tax.table <- data.frame(cbind(input.bold.lab.sheet,input.bold.taxonomy$Phylum,input.bold.taxonomy$Class,input.bold.taxonomy$Order))
  input.tax.table <- input.tax.table[,c("Process.ID", "Sample.ID","input.bold.taxonomy.Phylum","input.bold.taxonomy.Class","input.bold.taxonomy.Order")]
  names(input.tax.table) <- c("PID","SID","Phylum","Class","Order")
  PID.SID.tax.table <- data.frame("Sample" = c(input.tax.table$PID,input.tax.table$SID),
                                  "BOLD.Phylum" = c(input.tax.table$Phylum, input.tax.table$Phylum),
                                  "BOLD.Class" = c(input.tax.table$Class, input.tax.table$Class),
                                  "BOLD.Order" = c(input.tax.table$Order, input.tax.table$Order))
  PID.SID.tax.table$BOLD.Phylum[is.na(PID.SID.tax.table$BOLD.Phylum)] <- "unknown"
  PID.SID.tax.table$BOLD.Class[is.na(PID.SID.tax.table$BOLD.Class)] <- "unknown"
  PID.SID.tax.table$BOLD.Order[is.na(PID.SID.tax.table$BOLD.Order)] <- "unknown"
  
  #add Phylum, Class, and Order ID to contig table
  output <- merge(output, PID.SID.tax.table, by = "Sample")

  #flag contigs that have taxonomic mismatches (unknowns below Class do not count as mismatches)
  output$Phylum_Match <- ifelse(output$Phylum == output$BOLD.Phylum | output$BOLD.Phylum == "unknown" | is.na(output$Phylum) | is.na(output$BOLD.Phylum), "OK", "MISMATCH")
  output$Class_Match <- ifelse(output$Class == output$BOLD.Class | output$BOLD.Class == "unknown" | is.na(output$Class) | is.na(output$Class), "OK", "MISMATCH")
  output$Order_Match <- ifelse(output$Order == output$BOLD.Order | output$Order == "unknown" | output$BOLD.Order == "unknown" | is.na(output$Order) | is.na(output$Order), "OK", "MISMATCH")
  output$Tax_Match <- ifelse(output$Phylum_Match == "MISMATCH" | output$Class_Match == "MISMATCH" | output$Order_Match == "MISMATCH", "MISMATCH", "OK")
  
  #output table
  write.table(output, sprintf("%s_TaxonomicAssignments_AllContigs.tsv", plateid), append = F, quote = F, row.names = F, sep = "\t")
  
  #filter out mismatches
  output <- output[output$Tax_Match != "MISMATCH",]
}

#extract dominant contigs and output
df <- output[,c("Sample","ContigName","ReadCount","Sequence")]
df <- df[order(df$Sample,-df$ReadCount),]
df <- df[!duplicated(df$Sample),]
output.fasta <- DNAStringSet(df$Sequence)
names(output.fasta) <- paste(df$Sample,df$Contig,paste("Reads-",df$ReadCount, sep = ""), sep = "|")
writeXStringSet(output.fasta,sprintf("%s_DominantContigs.fasta",plateid))

#output contig table for dominant contigs only
output.dominants <- output[order(output$Sample,-output$ReadCount),]
output.dominants <- output.dominants[!duplicated(output.dominants$Sample),]
write.table(output.dominants, sprintf("%s_TaxonomicAssignments_DominantContigs.tsv", plateid), append = F, quote = F, row.names = F, sep = "\t")
```
