## Supplemental File 1 for "Barcode 100K Specimens: In a Single Nanopore Run": generate_histograms.R

```
#get arguments from bash script (plate map and contig FASTA file)
args <- commandArgs(trailingOnly = TRUE)
plateid <- args[1]
nbdemuxstep <- args[2]

#set working directory based on plate's folder, load required libraries
setwd(paste("/home/innovation-admin/DATA_INPUT/",plateid,"/", sep = ""))
library(Biostrings)
library(ggplot2)
library(cowplot)

###### READ COUNT HISTOGRAM ########
df <- read.table(paste(plateid,"_readcounts.txt", sep = ""), sep = "\t")
stages <- sapply(strsplit(df$V1, ":"), function(x) x[1])
counts <- sapply(strsplit(df$V1, ":"), function(x) x[2])
df$V1 <- as.character(stages)
df$V2 <- counts

#remove rows that were not applicable for this run
if (nbdemuxstep == "yes"){
  df <- df[df$V1 != "Input reads" & df$V1 != "After NB demux",]
}
df <- df[df$V2 != "NA",]
df$V2 <- as.numeric(as.character(df$V2))

#reformat step names
df$V1 <- gsub(" ","\n", df$V1)

#calculate read loss at each step and add colour based on percentage of reads that are left
total_rows <- nrow(df)

for (i in 1:total_rows) {
  if (i == 1) {
    df$V3[i] <- paste0("(", 100, "% remaining)")
    df$V4[i] <- paste0("= ", 0,"% drop")
    df$V5[i] <- 0
    df$V6[1] <- "#66BB6A"
  } else {
    df$V3[i] <- paste0("(", round(df$V2[i] / df$V2[1] * 100, digits = 0), "% remaining)")
    df$V4[i] <- paste0("= ", 100 - round(df$V2[i]/df$V2[i-1]*100, digits = 0), "% drop")
    df$V5[i] <- 100 - round(df$V2[i] / df$V2[i-1] * 100, digits = 0)
    if(df$V1[i] == "After\nbasecalling"){
      df$V6[i] <- ifelse(df$V5[i] <= 5, "#66BB6A",
                       ifelse(df$V5[i] > 5 && df$V5[i] <=10, "#FFB300", "firebrick" ))
    }
    if(df$V1[i] == "After\nNB\ndemux"){
      df$V6[i] <- ifelse(df$V5[i] <= 5, "#66BB6A",
                         ifelse(df$V5[i] > 6 && df$V5[i] <=10, "#FFB300", "firebrick" ))
    }
    if(df$V1[i] == "Assigned\nto\nplate/sample"){
      df$V6[i] <- ifelse(df$V5[i] <= 5, "#66BB6A",
                         ifelse(df$V5[i] > 6 && df$V5[i] <=10, "#FFB300", "firebrick" ))
    }
    if(df$V1[i] == "After\nsize\nfiltering"){
      df$V6[i] <- ifelse(df$V5[i] <= 15, "#66BB6A",
                         ifelse(df$V5[i] > 15 && df$V5[i] <=20, "#FFB300", "firebrick" ))
    }
    if(df$V1[i] == "After\nUMI\ndemux"){
      df$V6[i] <- ifelse(df$V5[i] <= 55, "#66BB6A",
                         ifelse(df$V5[i] > 55 && df$V5[i] <=60, "#FFB300", "firebrick" ))
    }
    if(df$V1[i] == "After\nprimer\ntrim"){
      df$V6[i] <- ifelse(df$V5[i] <= 5, "#66BB6A",
                         ifelse(df$V5[i] > 5 && df$V5[i] <=8, "#FFB300", "firebrick" ))
    }
    if(df$V1[i] == "Reads\nin\ncontigs"){
      df$V6[i] <- ifelse(df$V5[i] <= 30, "#66BB6A",
                         ifelse(df$V5[i] > 30 && df$V5[i] <=40, "#FFB300", "firebrick" ))
    }
    if(df$V1[i] == "Reads\nin\ndominants"){
      df$V6[i] <- ifelse(df$V5[i] <= 5, "#66BB6A",
                         ifelse(df$V5[i] > 5 && df$V5[i] <=10, "#FFB300", "firebrick" ))
    }
  }
}

#plot read loss histogram
plot1 <- ggplot(df, aes(y = V2, x = factor(df$V1, levels = df$V1), fill = V6)) +
  geom_bar(stat = "identity") +
  theme_bw() +
  theme(axis.text.x = element_text(size = 10, angle = 0, hjust = 0.5, color = "black", face = "bold"),
        axis.text.y = element_blank(),
        axis.title = element_blank(),
        panel.grid.major = element_blank(),
        panel.grid.minor = element_blank(),
        axis.ticks = element_blank()) +
  scale_y_continuous(expand = c(0, 0)) +
  scale_fill_identity() +
  geom_text(aes(label=paste(format(V2, big.mark = ",", trim = TRUE), V4, V3, "",sep = "\n"), fontface = "bold"),
            vjust=0,
            cex = 3) +
  coord_cartesian(ylim = c(0,max(df$V2) *1.15))
  

######## reads and contigs per specimen histograms ###########
platemap <- read.table(sprintf("/home/innovation-admin/DATA_INPUT/mapping_%s.txt",plateid), header = TRUE)
contigfasta <- readDNAStringSet(sprintf("/home/innovation-admin/DATA_INPUT/%s/%s_AllContigs.fasta",plateid,plateid))

#create master data frame from plate map and contig FASTA headers
df2 <- data.frame("Sample" = platemap$Label)
header <- names(contigfasta)
temp.df <- data.frame("Sample" = sapply(strsplit(header,"\\|"), function(x) x[1]),
                         "Contigs" = sapply(strsplit(header,"\\|"), function(x) x[2]),
                         "Reads" = sapply(strsplit(header,"\\|"), function(x) x[3]))
temp.df$Reads <- as.numeric(as.character(gsub("reads-","",temp.df$Reads)))
temp.reads <- aggregate(Reads ~ Sample, temp.df, sum)
temp.contigs <- aggregate(Contigs ~ Sample, temp.df, length)
df2 <- merge(df2, temp.reads, by = "Sample", all.x = TRUE)
df2 <- merge(df2, temp.contigs, by = "Sample", all.x = TRUE)
df2 <- replace(df2, is.na(df2), 0)

#subset data based on specific bin breaks for reads and contigs
df2$bin.reads = cut(df2$Reads, breaks = c(-1,0,5,10,15,20,25,50,100,200,300,400,500, Inf))
df2.reads <- data.frame(table(df2$bin.reads))
df2.reads$Var1 <- c("0","1-5","6-10","11-15","16-20","21-25","26-50","51-100","101-200","201-300","301-400","401-500",">500")

df2$bin.contigs = cut(df2$Contigs, breaks = c(-1,0,1,2,3,4,5,6,7,8,9,10, Inf))
df2.contigs <- data.frame(table(df2$bin.contigs))
df2.contigs$Var1 <- c("0","1","2","3","4","5","6","7","8","9","10",">10")

#plot histograms
plot2 <- ggplot(data = df2.reads) +
  geom_bar(stat = "identity", mapping = aes(x = factor(Var1, levels = c("0","1-5","6-10","11-15","16-20","21-25","26-50","51-100","101-200","201-300","301-400","401-500",">500")),
                                            y = Freq), fill = "blue", colour = "black") + 
  theme_bw() +
  ylab("No. Samples") +
  xlab("Reads/Sample") +
  theme(panel.grid.major.x = element_blank(),
        panel.grid.minor = element_blank(),
        axis.text.x = element_text(angle = 0, face = "bold"),
        plot.title = element_text(hjust = 0.5))
plot2 <- plot2 + coord_cartesian(ylim = c(0,max(df2.reads$Freq)*1.1))

plot3 <- ggplot(data = df2.contigs) +
  geom_bar(stat = "identity", mapping = aes(x = factor(Var1, levels = c("0","1","2","3","4","5","6","7","8","9","10",">10")),
                                            y = Freq), fill = "forestgreen", colour = "black") + 
  theme_bw() +
  ylab("No. Samples") +
  xlab("Contigs/Sample") +
  theme(panel.grid.major.x = element_blank(),
        panel.grid.minor = element_blank(),
        axis.text.x = element_text(angle = 0, face = "bold"),
        plot.title = element_text(hjust = 0.5))
plot3 <- plot3 + coord_cartesian(ylim = c(0,max(df2.contigs$Freq)*1.1))

#if this was done on a single plate, output the three plots to one PDF
names(platemap)[3] <- "Sample"
df3 <- merge(df2,platemap, by = "Sample")
df3 <- df3[df3$Reads != 0,]
df3 <- data.frame(table(df3$Sample.Plate.ID))
names(df3) <- c("Plate", "Count")
df3$Colour <- ifelse(df3$Count >= 75, "#66BB6A",
                     ifelse(df3$Count < 75 & df3$Count >= 50, "#FFB300", "firebrick"))

if (nbdemuxstep == "yes" | length(df3$Plate) == 1){
  pdf(sprintf("%s_SummaryHistograms.pdf",plateid), width = 8.5, height = 11)
  print(ggdraw() +
    draw_plot(plot1, x = 0, y = 0.2, width = 1, height = 0.6) +
    draw_plot_label(label = sprintf("%s Summary Plots",plateid), size = 35, x = -0.35, y = 0.95))
  print(ggdraw() +
    draw_plot(plot2, x = 0, y = 0.505, width = 1, height = 0.45) +
    draw_plot(plot3, x = 0, y = 0.05, width = 1, height = 0.45))
  dev.off()
} else {
  #if this was done on more than one plate, plot success-per-plate histogram
  plot4 <- ggplot(df3, aes(y = Plate, x = Count, fill = Colour)) +
    geom_bar(stat = "identity") +
    theme_bw() +
    theme(axis.text.x = element_text(size = 10, angle = 0, hjust = 0.5, color = "black", face = "bold"),
          axis.text.y = element_text(size = 10, angle = 0, hjust = 0.5, color = "black", face = "bold"),
          axis.title.y = element_blank(),
          axis.title.x = element_text(face = "bold"),
          panel.grid.major.y = element_blank(),
          panel.grid.minor = element_blank(),
          axis.ticks = element_blank()) +
    scale_x_continuous(expand = c(0, 1)) +
    scale_fill_identity() +
    labs(x = "Sequence Count") +
    coord_cartesian(xlim = c(0,100))
  
  #plot all four plots on one PDF
  pdf(sprintf("%s_SummaryHistograms.pdf",plateid), width = 8.5, height = 11)
  
  print(ggdraw() +
    draw_plot(plot1, x = 0, y = 0.2, width = 1, height = 0.6) +
    draw_plot_label(label = sprintf("%s Summary Plots",plateid), size = 35, x = 0.5, y = 0.95, hjust = 0.5))
  
  print(ggdraw() +
    draw_plot(plot2, x = 0, y = 0.505, width = 1, height = 0.45) +
    draw_plot(plot3, x = 0, y = 0.05, width = 1, height = 0.45))
  
  if(length(df3$Plate) <= 24){
    plot4_height <- 0.35
    print(ggdraw() + draw_plot(plot4, x = 0, y = 1 - plot4_height, width = 1, height = plot4_height))
  }
  
  if(length(df3$Plate) > 24 & length(df3$Plate) <= 48){
    plot4_height <- 0.70
    print(ggdraw() + draw_plot(plot4, x = 0, y = 1 - plot4_height, width = 1, height = plot4_height))
  }
   
  if(length(df3$Plate) > 48 & length(df3$Plate) <= 72){
    plot4_height <- 0.85
    print(ggdraw() + draw_plot(plot4, x = 0, y = 1 - plot4_height, width = 1, height = plot4_height))
  }
  
  if(length(df3$Plate) > 72){
    plot4_height <- 1
    print(ggdraw() + draw_plot(plot4 + theme(axis.text.y = element_text(size = 5, angle = 0, hjust = 0.5, color = "black", face = "bold")), x = 0, y = 1 - plot4_height, width = 1, height = plot4_height))
  }
  dev.off()
}
```
