## Supplemental File 1 for "Barcode 100K Specimens: In a Single Nanopore Run": get_parameters.R

```
#set working directory and load required libraries
setwd("/home/innovation-admin/DATA_INPUT/")
library("readxl")

#input run info tab from parameters file
input.runinfo <- data.frame(read_excel(list.files(pattern = "parameters.*\\.xlsx"),sheet = "Run Info"))

#remove unused rows
input.runinfo <- input.runinfo[!is.na(input.runinfo$Run.Name),]

#output parameters text file
write.table(input.runinfo, "runinfo.txt", append = F, quote = F, row.names = F, sep = "\t")

#input UMI map based on user selection
args <- commandArgs(trailingOnly = TRUE)
umitype <- args[1]
nbdemuxstep <- args[2]

if (umitype == "sequel"){
  input.umitype <- data.frame(read_excel(list.files(pattern = "parameters.*\\.xlsx"),sheet = "UMI Map - Sequel"))
  
  if(nbdemuxstep == "no"){ #output all UMIs as one mapping file
    write.table(input.umitype, sprintf("mapping_%s.txt", input.runinfo$Library.Plate.Name[1]), append = F, quote = F, row.names = F, sep = "\t")
  }
  
  if(nbdemuxstep == "yes"){ #output one UMI map for each library
    uniqueplates <- unique(input.umitype$Sample.Plate.ID)
    for (f in uniqueplates) {
      platemap <- input.umitype[which(input.umitype$Sample.Plate.ID == f),]
      write.table(platemap, sprintf("mapping_%s.txt", f), append = F, quote = F, row.names = F, sep = "\t")
    }
  }
}

if (umitype == "asym"){
  input.umitype <- data.frame(read_excel(list.files(pattern = "parameters.*\\.xlsx"),sheet = "UMI Map - Asymmetrical"))
  
  if(nbdemuxstep == "no"){ #output all UMIs as one mapping file
    write.table(input.umitype, sprintf("mapping_%s.txt", input.runinfo$Library.Plate.Name[1]), append = F, quote = F, row.names = F, sep = "\t")
  }
  
  if(nbdemuxstep == "yes"){ #output one UMI map for each library
    uniqueplates <- unique(input.umitype$Sample.Plate.ID)
    for (f in uniqueplates) {
      platemap <- input.umitype[which(input.umitype$Sample.Plate.ID == f),]
      write.table(platemap, sprintf("mapping_%s.txt", f), append = F, quote = F, row.names = F, sep = "\t")
    }
  }
  
}

if (umitype == "sym"){
  input.umitype <- data.frame(read_excel(list.files(pattern = "parameters.*\\.xlsx"),sheet = "UMI Map - Symmetrical"))
  
  if(nbdemuxstep == "no"){ #output all UMIs as one mapping file
    write.table(input.umitype, sprintf("mapping_%s.txt", input.runinfo$Library.Plate.Name[1]), append = F, quote = F, row.names = F, sep = "\t")
  }
  
  if(nbdemuxstep == "yes"){ #output one UMI map for each plate
    uniqueplates <- unique(input.umitype$Sample.Plate.ID)
    for (f in uniqueplates) {
      platemap <- input.umitype[which(input.umitype$Sample.Plate.ID == f),]
      write.table(platemap, sprintf("mapping_%s.txt", f), append = F, quote = F, row.names = F, sep = "\t")
    }
  }
}
```
