## Supplemental File 1 for "Barcode 100K Specimens: In a Single Nanopore Run": ONT.sh

```
#!/bin/bash

#set number of cores to use
cores=24

cd /home/innovation-admin/DATA_INPUT/

clear
echo -e "\e[1m********** ONT Data Processing - CBG Innovation Unit **********\e[0m\n"
echo -e "This will process ONT data in various ways depending on the user's selected criteria.\n"
echo -e "\e[1mRequirements:\e[0m\n"
echo -e "1) Starting data files (see below for details), placed in DATA_INPUT.\n"
echo -e "2) One parameters file (parameters_XXXXX.xlsx), placed in DATA_INPUT.\n"
echo -e "3) OPTIONAL: BOLD datasheet file including the LAB SHEET & TAXONOMY tabs. Should be named taxonomy_XXXXX.xlsx and placed in DATA_INPUT.\n\n"

echo -e "\e[1mIf starting from...\e[0m\n"
echo -e "\e[1mRaw ONT data\e[0m: Put POD5 files in DATA_INPUT.\n"
echo -e "\e[1mAlready basecalled FASTQ/FASTA file(s)\e[0m: Put all files in DATA_INPUT.\n\n"

echo "Press any key to continue..."
read -n 1 -s

#define the menu options
options=(
  1 "Basecall" on
  2 "Native barcode demultiplex" off
  3 "Length filtering" on
  4 "UMI demultiplex" on
  5 "Primer trim" on
  6 "Contig assembly" on
)

#display the checklist and save the selected options to a variable
selected=$(dialog --stdout --checklist "Select options using <space> then press <enter> to continue:" 0 0 0 "${options[@]}")

#if UMI demux is selected, query user for type of UMIs
if [[ $selected == *"4"* ]]; then
	umioptions=(
  	1 "Asymmetrical UMIs (96 x 96)" off
  	2 "Asymmetrical UMIs (96 x 24)" on
  	3 "Symmetrical UMIs (96 x 1)" off
	)
	umiselected=$(dialog --stdout --radiolist "Select UMI type using <space> then press <enter> to continue:" 0 0 0 "${umioptions[@]}")
fi

#print the selected options and promt user to confirm
clear
echo "You have selected the following steps:"
for option in $selected; do
  case $option in
    1) echo "- Basecall";;
    2) echo "- Native barcode demultiplex";;
	3) echo "- Length filtering";;
    4) echo "- UMI demultiplex";;
	5) echo "- Primer trim";;
	6) echo "- Contig assembly";;
  esac
done
echo -e "\n\n"

if [[ $selected == *"4"* ]]; then
	echo "You have selected the following UMIs:"
	for umioption in $umiselected; do
	  case $umioption in
	    1) echo "- Asymmetrical UMIs (96 x 96)";;
	    2) echo "- Asymmetrical UMIs (96 x 24)";;
	    3) echo "- Symmetrical UMIs (96 x 1)";;
	  esac
	done
	echo -e "\n\n"
fi

read -p "Do you want to proceed? Type 'yes' to confirm: " choice

#proceed with script if user typed 'yes'
case $choice in
  yes)
	echo "Script running..."
    exec &> /home/innovation-admin/DATA_INPUT/terminalhistory.log #redirect terminal history to log file
    echo "Script started at $(date +"%Y-%m-%d %H:%M:%S")"
    ;;
  *)
    echo "Aborting."
    exit 1
    ;;
esac

#assign default values to read count variables
inputreadcount=NA
basecallreadcount=NA
nbdemuxreadcount=NA
plateassignreadcount=NA
sizefilterreadcount=NA
umidemuxreadcount=NA
ptrimreadcount=NA
contigreadcount=NA
dominantreadcount=NA

#assign user options to variables
if [[ $selected == *"1"* ]]; then
	basecallstep="yes"
else
	basecallstep="no"
fi

if [[ $selected == *"2"* ]]; then
	nbdemuxstep="yes"
else
	nbdemuxstep="no"
fi

if [[ $selected == *"3"* ]]; then
	sizefilterstep="yes"
else
	sizefilterstep="no"
fi

if [[ $selected == *"4"* ]]; then
	umidemuxstep="yes"
else
	umidemuxstep="no"
fi

if [[ $selected == *"5"* ]]; then
	ptrimstep="yes"
	ptrimreadcount=0
else
	ptrimstep="no"
fi

if [[ $selected == *"6"* ]]; then
	contigstep="yes"
	contigreadcount=0
	dominantreadcount=0
else
	contigstep="no"
fi

umitype="none" #default if UMI demux is not requested, but will be updated below if requested
if [[ $umiselected == *"1"* ]]; then
	umitype="sequel"
fi

if [[ $umiselected == *"2"* ]]; then
	umitype="asym"
fi

if [[ $umiselected == *"3"* ]]; then
	umitype="sym"
fi

#collect and reformat parameters information
Rscript /home/innovation-admin/SCRIPTS/SUBSCRIPTS/get_parameters.R $umitype $nbdemuxstep

#check to see if BOLD taxonomy file is present
boldtaxcount=$(ls taxonomy*.xlsx | wc -l)
if [[ "$boldtaxcount" -eq 1 ]]; then
    boldtaxpresent="yes"
else
    boldtaxpresent="no"
fi

#check to see if FASTQ file is present and consolidate into single file if true and convert to FASTA
fastqcount=$(ls *.fastq | wc -l)
if [[ "$fastqcount" -gt 0 ]]; then
    fastqpresent="yes"
else
    fastqpresent="no"
fi

#check to see if FASTA file is present and consolidate into single file if true
fastacount=$(ls *.fasta | wc -l)
if [[ "$fastacount" -gt 0 ]]; then
    fastapresent="yes"
else
    fastapresent="no"
fi

#check to see if pod5 files are present
podcount=$(ls *.pod5 | wc -l)
if [[ "$podcount" -gt 0 ]]; then
    podpresent="yes"
else
    podpresent="no"
fi

#if no input files present, exit script
if [[ "$fastqpresent" == "no" && "$fastapresent" == "no" && "$podpresent" == "no" ]]; then
	echo "ERROR: NO INPUT FILES DETECTED!"
	exit
fi

#if FASTQ or FASTA present but basecalling requested, exit script
if [[ $$basecallstep == "yes" && $podpresent == "no" ]]; then
 	echo "ERROR: BASECALLING REQUIRES POD5 FILES."
 	exit
fi

#if FASTA present but NB demux requested, exit script
if [[ nbdemuxstep == "yes" && $fastapresent == "yes" ]]; then
	echo "ERROR: ONLY FASTQ FILES CAN BE DEMULTIPLEXED WITH NATIVE BARCODES."
	exit
fi

#Extract barcode list if NB demux requested, or plate ID if NB demux not requested
if [[ $nbdemuxstep == "yes" ]]; then
	barcodes=$(tail -n +2 runinfo.txt | cut -f 3 -s)
fi

if [[ $nbdemuxstep == "no" ]]; then
	plateid=$(tail -n +2 runinfo.txt | cut -f 2 -s)
fi

########################################################################################################################################################################
# basecall
########################################################################################################################################################################

#if basecalling was requested, run dorado
if [[ $basecallstep == "yes" ]]; then
	echo -e "\n****************** Basecalling data...\n"
	#exec &>/dev/tty #redirect terminal history to terminal so basecalling progress can be observed
	dorado basecaller /opt/ont/dorado/data/dna_r10.4.1_e8.2_400bps_sup@v4.2.0 ./ --emit-fastq --device cuda:0 > all.fastq
	#exec &> /home/ccdb/DATA_INPUT/terminalhistory.log #redirect terminal history back to log file
	rm *.pod5
	basecallreadcount=$(echo $(cat all.fastq|wc -l)/4|bc)
fi

#if this is the last step, clean up data and exit script
if [[ $basecallstep == "yes" && $nbdemuxstep == "no" && $sizefilterstep == "no" && $umidemuxstep == "no" && $ptrimstep == "no" && $contigstep == "no" ]]; then
	if [[ $fastqpresent == "yes" ]]; then
		paste - - - - < all.fastq | cut -f 1,2 | sed 's/^@/>/' | tr "\t" "\n" > all.fasta #convert to FASTA
	fi
	echo -e "\n Basecalling complete."
	exit
fi

########################################################################################################################################################################
# native barcoding demultiplexing
########################################################################################################################################################################

#if this is the first step, concatenate all input FASTQ files into a single file if needed for UMI demux
if [[ $basecallstep == "no" && $nbdemuxstep == "yes" && $umidemuxstep == "yes" ]]; then
	cat *.fastq > all.fastq2
	rm *.fastq
	mv all.fastq2 all.fastq
	inputreadcount=$(echo $(cat all.fastq|wc -l)/4|bc)
fi

#if NB demultiplexing was requested, demultiplex native barcodes using Dorado
if [[ $nbdemuxstep == "yes" ]]; then
	
	#set nbdemuxreadcount variable to 0
	nbdemuxreadcount=0

	#demultiplex native barcodes using Dorado
	echo -e "\n****************** Demultiplexing native barcodes...\n"
	dorado demux --kit-name SQK-NBD114-24 --output-dir ./BARCODED --emit-fastq all.fastq 
	rm all.fastq

	#move into BARCODED subdirectory
	cd BARCODED
	
	#remove unclassified reads
	rm unclassified.fastq

	#remove NB barcode kit name from demuxed FASTQ files
	rename 's/SQK-NBD114-24_//g' *.fastq

	#create subfolders based on barcode names, then move FASTQ files into them and rename to all.fastq and convert to FASTA
	for file in barcode*.fastq; do
	    #extract barcode name from file name
	    barcodename="$(basename "$file" .fastq)"
	     
	    #create a directory with the barcode name
	    mkdir -m 777 "$barcodename"
	      
	    #move the corresponding file to the directory
	    mv "$file" "$barcodename/"
	      
	    #rename FASTQ file to plate/sample name
	    cd $barcodename
	    mv $file all.fastq
	    tempreadcount=$(echo $(cat all.fastq | wc -l)/4 | bc)
		nbdemuxreadcount=$((nbdemuxreadcount + tempreadcount))
		paste - - - - < all.fastq | cut -f 1,2 | sed 's/^@/>/' | tr "\t" "\n" > all.fasta
		rm all.fastq
      	cd ../
 	done

 	#move back into working directory
 	cd /home/innovation-admin/DATA_INPUT

 	#move barcoded folders to working directory
	for barcode in $barcodes
	do
	  mv "/home/innovation-admin/DATA_INPUT/BARCODED/$barcode" /home/innovation-admin/DATA_INPUT
	done

	#rename barcoded folders by plate specified in runinfo.txt
	while IFS=$'\t' read -r _ new_name old_name _ _ _ _ _ _
	do
	  #Construct the paths to the old and new folders
	  old_path="/home/innovation-admin/DATA_INPUT/$old_name"
	  new_path="/home/innovation-admin/DATA_INPUT/$new_name"

	  #Rename the folder if it exists
	  if [ -d "$old_path" ]
	  then
	    mv "$old_path" "$new_path"
	  fi
	done < runinfo.txt

	#process the last line of the input file
	read -r _ new_name old_name _ _ _ _ _ _ <<< $(tail -n 1 runinfo.txt)
	old_path="/home/innovation-admin/DATA_INPUT/$old_name"
	new_path="/home/innovation-admin/DATA_INPUT/$new_name"
	if [ -d "$old_path" ]
	then
	  mv "$old_path" "$new_path"
	fi

	#delete BARCODED folder
	rm -r ./BARCODED
fi

#if NB demultiplexing is the last step, clean up data then exit script
if [[ $nbdemuxstep == "yes" && $umidemuxstep == "no" && $sizefilterstep == "no" && $ptrimstep == "no" && $contigstep == "no" ]]; then
	mkdir -m 777 NB_demultiplexed_files_FASTQ
	mkdir -m 777 NB_demultiplexed_files_FASTA
	for dir in /home/innovation-admin/DATA_INPUT/*/; do 
    if [[ "$dir" != "/home/innovation-admin/DATA_INPUT/NB_demultiplexed_files_FASTQ/" && "$dir" != "/home/innovation-admin/DATA_INPUT/NB_demultiplexed_files_FASTA/" ]]; then
	    cd $dir
	    plateid=$(basename "$PWD")
	    paste - - - - < all.fastq | cut -f 1,2 | sed 's/^@/>/' | tr "\t" "\n" > "$plateid".fasta #convert to FASTA
	    mv all.fastq "$plateid".fastq
	    mv "$plateid".fastq "/home/innovation-admin/DATA_INPUT/NB_demultiplexed_files_FASTQ"
	    mv "$plateid".fasta "/home/innovation-admin/DATA_INPUT/NB_demultiplexed_files_FASTA"
	    cd ../
	    rm -r $dir
	  fi
	done
	echo -e "\n Native baracoding demultiplexing complete."
	exit
fi


#if NB demultiplexing was not requested, reformat input data depending on whether all input files need to be kept separate or merged into single file for UMI demux
if [[ $nbdemuxstep == "no" ]]; then
	if [[ $basecallstep == "yes" ]]; then #file was generated via basecalling and does not require merging, just convert all.fastq to FASTA and move into plate folder
		mkdir -m 777 /home/innovation-admin/DATA_INPUT/"$plateid"
		mv all.fastq /home/innovation-admin/DATA_INPUT/"$plateid"
		paste - - - - < /home/innovation-admin/DATA_INPUT/"$plateid"/all.fastq | cut -f 1,2 | sed 's/^@/>/' | tr "\t" "\n" > /home/innovation-admin/DATA_INPUT/"$plateid"/all.fasta 
	fi

	if [[ $basecallstep == "no" && $umidemuxstep == "no" && $fastqpresent == "yes" ]]; then #file(s) were supplied by user and do not require merging, just convert all to FASTA and move into plate folder
		mkdir -m 777 /home/innovation-admin/DATA_INPUT/"$plateid"
		for f in *.fastq; do
			paste - - - - < $f | cut -f 1,2 | sed 's/^@/>/' | tr "\t" "\n" > $f.fasta
			rename 's/.fastq.fasta/.fasta/g' $f.fasta
			rm $f
		done
		mv *.fasta /home/innovation-admin/DATA_INPUT/"$plateid"
	fi

	if [[ $basecallstep == "no" && $umidemuxstep == "no" && $fastapresent == "yes" ]]; then #file(s) were supplied by user and do not require merging, just move into plate folder
		mkdir -m 777 /home/innovation-admin/DATA_INPUT/"$plateid"
		mv *.fasta /home/innovation-admin/DATA_INPUT/"$plateid"
	fi

	if [[ $basecallstep == "no" && $umidemuxstep == "yes" && $fastqpresent == "yes" ]]; then #files require merging, concatenate all FASTQ files into all.fastq, then convert to all.fasta
		cat *.fastq > all.fastq2
		rm *.fastq
		mv all.fastq2 all.fastq
		mkdir -m 777 /home/innovation-admin/DATA_INPUT/"$plateid"
		mv all.fastq /home/innovation-admin/DATA_INPUT/"$plateid"
		paste - - - - < /home/innovation-admin/DATA_INPUT/"$plateid"/all.fastq | cut -f 1,2 | sed 's/^@/>/' | tr "\t" "\n" > /home/innovation-admin/DATA_INPUT/"$plateid"/all.fasta #convert to FASTA
		rm all.fastq
		inputreadcount=$(grep -c ">" /home/innovation-admin/DATA_INPUT/"$plateid"/all.fasta)
	fi

	if [[ $basecallstep == "no" && $umidemuxstep == "yes" && $fastapresent == "yes" ]]; then #files require merging, concatenate all FASTA files into all.fasta
		cat *.fasta > all.fasta2
		rm *.fasta
		mv all.fasta2 all.fasta
		mkdir -m 777 /home/innovation-admin/DATA_INPUT/"$plateid"
		mv all.fastq /home/innovation-admin/DATA_INPUT/"$plateid"
		paste - - - - < /home/innovation-admin/DATA_INPUT/"$plateid"/all.fastq | cut -f 1,2 | sed 's/^@/>/' | tr "\t" "\n" > /home/innovation-admin/DATA_INPUT/"$plateid"/all.fasta #convert to FASTA
		inputreadcount=$(grep -c ">" /home/innovation-admin/DATA_INPUT/"$plateid"/all.fasta)
	fi

fi


# <<<<<<<<<<< BY THIS POINT, ALL DATA SHOULD BE IN FOLDERS NAMED AFTER PLATE(S) AND BE A EITHER SINGLE FILE NAMED all.fasta (NEEDS UMI DEMUX) OR MULTIPLE FASTA FILES (DOES NOT NEED UMI DEMUX)


#Process each plate/sample folder seperately for subsequent steps
platetask(){
  
 	#move into the first plate/sample folder
  	cd "$dir"
	
  	#extract plate/sample-specific parameters from runinfo.txt file
	plateid=$(basename "$PWD")
	platedata=($(grep "$plateid" /home/innovation-admin/DATA_INPUT/runinfo.txt))
	runid=${platedata[0]}
	forprimername=${platedata[3]}
	revprimername=${platedata[4]}
	minreadlen=${platedata[5]}
	maxreadlen=${platedata[6]}
	ampsize=${platedata[7]}
	minreads=${platedata[8]}
	trimlength=$(awk "BEGIN { printf \"%.0f\", $ampsize * 0.15 }")
	reflib=${platedata[9]}
	sintax=${platedata[10]}

	#create temporary FASTA file for forward primers
	for primer in $(echo "$forprimername" | tr ',' '\n'); do
	    if grep -A 1 "$primer" /home/innovation-admin/PRIMERS/primerDB.fasta; then
	        grep -A 1 "$primer" /home/innovation-admin/PRIMERS/primerDB.fasta >> temp_fwd_primers.fa
	    else
	        echo -e "ERROR: $primer not found in file PrimerDB.fasta.\n"
	        rm temp_fwd_primers.fa
	        exit
	    fi
	done

	#reverse-complement forward primers
	seqtk seq -r -c temp_fwd_primers.fa > temp_fwd_primers_rc.fa

	#create temporary FASTA file for reverse primers
	for primer in $(echo "$revprimername" | tr ',' '\n'); do
	    if grep -A 1 "$primer" /home/innovation-admin/PRIMERS/primerDB.fasta; then
	        grep -A 1 "$primer" /home/innovation-admin/PRIMERS/primerDB.fasta >> temp_rev_primers.fa
	    else
	        echo -e "ERROR: $primer not found in file PrimerDB.fasta.\n"
	        rm temp_rev_primers.fa
	        exit
	    fi
	done

	#reverse-complement reverse primers
	seqtk seq -r -c temp_rev_primers.fa > temp_rev_primers_rc.fa

	#make directories to hold original reads and contig read FASTA files (if necessary)
	mkdir -m 777 /home/innovation-admin/DATA_INPUT/"$plateid"/Original_Reads
	
	if [[ $contigstep == "yes" ]]; then
		mkdir -m 777 /home/innovation-admin/DATA_INPUT/"$plateid"/Contig_Component_Reads
		mkdir -m 777 /home/innovation-admin/DATA_INPUT/"$plateid"/Contig_Consensus_Sequences
	fi

	#count reads assigned to plate/sample
	if [[ $umidemuxstep == "yes" ]]; then
		plateassignreadcount=$(grep -c ">" all.fasta)
	fi

	########################################################################################################################################################################
	# size filtering
	########################################################################################################################################################################

	if [[ $sizefilterstep == "yes" && $umidemuxstep == "yes" ]]; then
		echo -e "\n****************** Size filtering $plateid...\n"
		
		#filter raw reads by min and max length (split into files of 1M reads if necessary)
		split -l 2000000 all.fasta part_
		for file in part_*; do
		    cutadapt -j $((cores-1)) -m $minreadlen -M $maxreadlen -o filtered_${file} ${file}&
		done
		wait
		cat filtered_part_* > all.filtered.fasta
		rm part_*
		rm filtered_part_*
		rm all.fasta
		mv all.filtered.fasta all.fasta

		#get read count of size filtered reads
		sizefilterreadcount=$(grep -c "^>" all.fasta)
	fi

	if [[ $sizefilterstep == "yes" && $umidemuxstep == "no" ]]; then
		echo -e "\n****************** Size filtering samples...\n"
		
		#filter raw reads by min and max length and count filtered reads
		sizefilterreadcount=0
		for file in *.fasta; do
		    cutadapt -j $((cores-1)) -m $minreadlen -M $maxreadlen -o "${file}_filtered" "$file"
		    mv "$file" Original_Reads
		    mv "${file}_filtered" "$file"
		    reads=$(grep -c "^>" "$file")
		    sizefilterreadcount=$((sizefilterreadcount + reads))
		done
	fi

	#if size filtering is the last step, continue to next plate/sample (if exists), otherwise skip to the end of the plate loop
	if [[ $sizefilterstep == "yes" && $umidemuxstep == "no" && $ptrimstep == "no" && $contigstep == "no" ]]; then
		return
	fi

	########################################################################################################################################################################
	# UMI demultiplex
	########################################################################################################################################################################

	#if UMI demux requested, demultiplex all.fasta into individual sample files based on umi type provided
	if [[ $umidemuxstep == "yes" ]]; then

		##### asymmetrical UMIs (96 x 24)
		if [[ $umitype == "asym" ]]; then

			echo -e "\n****************** Demultiplexing 96X24 UMIs for $plateid...\n"
			#demultiplex (allow up to 2 mismatches and indels)
			cutadapt -j $((cores-1)) -e 2 -O 12 -g file:/home/innovation-admin/PRIMERS/PacBio_V2_UMI1-96.fasta -o "trimmed-{name}.for5.fasta" all.fasta #search for all 96 UMIs at 5' end
			seqtk seq -r trimmed-unknown.for5.fasta > trimmed-unknown.for5.rc.fasta #rev-comp reads that don't have 5'UMI
			cutadapt -j $((cores-1)) -e 2 -O 12 -g file:/home/innovation-admin/PRIMERS/PacBio_V2_UMI1-96.fasta -o "trimmed-{name}.for3.fasta" trimmed-unknown.for5.rc.fasta #search for all 96 UMIs at new 5' end
			rm *unknown*.fasta all.fasta
			
			#merge forward and reverse split files into a single fasta file
			cat trimmed-bc1001.for5.fasta trimmed-bc1001.for3.fasta > bc1001.fasta
			cat trimmed-bc1002.for5.fasta trimmed-bc1002.for3.fasta > bc1002.fasta
			cat trimmed-bc1003.for5.fasta trimmed-bc1003.for3.fasta > bc1003.fasta
			cat trimmed-bc1004.for5.fasta trimmed-bc1004.for3.fasta > bc1004.fasta
			cat trimmed-bc1005.for5.fasta trimmed-bc1005.for3.fasta > bc1005.fasta
			cat trimmed-bc1006.for5.fasta trimmed-bc1006.for3.fasta > bc1006.fasta
			cat trimmed-bc1007.for5.fasta trimmed-bc1007.for3.fasta > bc1007.fasta
			cat trimmed-bc1008.for5.fasta trimmed-bc1008.for3.fasta > bc1008.fasta
			cat trimmed-bc1009.for5.fasta trimmed-bc1009.for3.fasta > bc1009.fasta
			cat trimmed-bc1010.for5.fasta trimmed-bc1010.for3.fasta > bc1010.fasta
			cat trimmed-bc1011.for5.fasta trimmed-bc1011.for3.fasta > bc1011.fasta
			cat trimmed-bc1012.for5.fasta trimmed-bc1012.for3.fasta > bc1012.fasta
			cat trimmed-bc1013.for5.fasta trimmed-bc1013.for3.fasta > bc1013.fasta
			cat trimmed-bc1014.for5.fasta trimmed-bc1014.for3.fasta > bc1014.fasta
			cat trimmed-bc1015.for5.fasta trimmed-bc1015.for3.fasta > bc1015.fasta
			cat trimmed-bc1016.for5.fasta trimmed-bc1016.for3.fasta > bc1016.fasta
			cat trimmed-bc1017.for5.fasta trimmed-bc1017.for3.fasta > bc1017.fasta
			cat trimmed-bc1018.for5.fasta trimmed-bc1018.for3.fasta > bc1018.fasta
			cat trimmed-bc1019.for5.fasta trimmed-bc1019.for3.fasta > bc1019.fasta
			cat trimmed-bc1020.for5.fasta trimmed-bc1020.for3.fasta > bc1020.fasta
			cat trimmed-bc1021.for5.fasta trimmed-bc1021.for3.fasta > bc1021.fasta
			cat trimmed-bc1022.for5.fasta trimmed-bc1022.for3.fasta > bc1022.fasta
			cat trimmed-bc1023.for5.fasta trimmed-bc1023.for3.fasta > bc1023.fasta
			cat trimmed-bc1024.for5.fasta trimmed-bc1024.for3.fasta > bc1024.fasta
			cat trimmed-bc1025.for5.fasta trimmed-bc1025.for3.fasta > bc1025.fasta
			cat trimmed-bc1026.for5.fasta trimmed-bc1026.for3.fasta > bc1026.fasta
			cat trimmed-bc1027.for5.fasta trimmed-bc1027.for3.fasta > bc1027.fasta
			cat trimmed-bc1028.for5.fasta trimmed-bc1028.for3.fasta > bc1028.fasta
			cat trimmed-bc1029.for5.fasta trimmed-bc1029.for3.fasta > bc1029.fasta
			cat trimmed-bc1030.for5.fasta trimmed-bc1030.for3.fasta > bc1030.fasta
			cat trimmed-bc1031.for5.fasta trimmed-bc1031.for3.fasta > bc1031.fasta
			cat trimmed-bc1032.for5.fasta trimmed-bc1032.for3.fasta > bc1032.fasta
			cat trimmed-bc1033.for5.fasta trimmed-bc1033.for3.fasta > bc1033.fasta
			cat trimmed-bc1034.for5.fasta trimmed-bc1034.for3.fasta > bc1034.fasta
			cat trimmed-bc1035.for5.fasta trimmed-bc1035.for3.fasta > bc1035.fasta
			cat trimmed-bc1036.for5.fasta trimmed-bc1036.for3.fasta > bc1036.fasta
			cat trimmed-bc1037.for5.fasta trimmed-bc1037.for3.fasta > bc1037.fasta
			cat trimmed-bc1038.for5.fasta trimmed-bc1038.for3.fasta > bc1038.fasta
			cat trimmed-bc1039.for5.fasta trimmed-bc1039.for3.fasta > bc1039.fasta
			cat trimmed-bc1040.for5.fasta trimmed-bc1040.for3.fasta > bc1040.fasta
			cat trimmed-bc1041.for5.fasta trimmed-bc1041.for3.fasta > bc1041.fasta
			cat trimmed-bc1042.for5.fasta trimmed-bc1042.for3.fasta > bc1042.fasta
			cat trimmed-bc1043.for5.fasta trimmed-bc1043.for3.fasta > bc1043.fasta
			cat trimmed-bc1044.for5.fasta trimmed-bc1044.for3.fasta > bc1044.fasta
			cat trimmed-bc1045.for5.fasta trimmed-bc1045.for3.fasta > bc1045.fasta
			cat trimmed-bc1046.for5.fasta trimmed-bc1046.for3.fasta > bc1046.fasta
			cat trimmed-bc1047.for5.fasta trimmed-bc1047.for3.fasta > bc1047.fasta
			cat trimmed-bc1048.for5.fasta trimmed-bc1048.for3.fasta > bc1048.fasta
			cat trimmed-bc1049.for5.fasta trimmed-bc1049.for3.fasta > bc1049.fasta
			cat trimmed-bc1050.for5.fasta trimmed-bc1050.for3.fasta > bc1050.fasta
			cat trimmed-bc1051.for5.fasta trimmed-bc1051.for3.fasta > bc1051.fasta
			cat trimmed-bc1052.for5.fasta trimmed-bc1052.for3.fasta > bc1052.fasta
			cat trimmed-bc1053.for5.fasta trimmed-bc1053.for3.fasta > bc1053.fasta
			cat trimmed-bc1054.for5.fasta trimmed-bc1054.for3.fasta > bc1054.fasta
			cat trimmed-bc1055.for5.fasta trimmed-bc1055.for3.fasta > bc1055.fasta
			cat trimmed-bc1056.for5.fasta trimmed-bc1056.for3.fasta > bc1056.fasta
			cat trimmed-bc1057.for5.fasta trimmed-bc1057.for3.fasta > bc1057.fasta
			cat trimmed-bc1058.for5.fasta trimmed-bc1058.for3.fasta > bc1058.fasta
			cat trimmed-bc1059.for5.fasta trimmed-bc1059.for3.fasta > bc1059.fasta
			cat trimmed-bc1060.for5.fasta trimmed-bc1060.for3.fasta > bc1060.fasta
			cat trimmed-bc1061.for5.fasta trimmed-bc1061.for3.fasta > bc1061.fasta
			cat trimmed-bc1062.for5.fasta trimmed-bc1062.for3.fasta > bc1062.fasta
			cat trimmed-bc1063.for5.fasta trimmed-bc1063.for3.fasta > bc1063.fasta
			cat trimmed-bc1064.for5.fasta trimmed-bc1064.for3.fasta > bc1064.fasta
			cat trimmed-bc1065.for5.fasta trimmed-bc1065.for3.fasta > bc1065.fasta
			cat trimmed-bc1066.for5.fasta trimmed-bc1066.for3.fasta > bc1066.fasta
			cat trimmed-bc1067.for5.fasta trimmed-bc1067.for3.fasta > bc1067.fasta
			cat trimmed-bc1068.for5.fasta trimmed-bc1068.for3.fasta > bc1068.fasta
			cat trimmed-bc1069.for5.fasta trimmed-bc1069.for3.fasta > bc1069.fasta
			cat trimmed-bc1070.for5.fasta trimmed-bc1070.for3.fasta > bc1070.fasta
			cat trimmed-bc1071.for5.fasta trimmed-bc1071.for3.fasta > bc1071.fasta
			cat trimmed-bc1072.for5.fasta trimmed-bc1072.for3.fasta > bc1072.fasta
			cat trimmed-bc1073.for5.fasta trimmed-bc1073.for3.fasta > bc1073.fasta
			cat trimmed-bc1074.for5.fasta trimmed-bc1074.for3.fasta > bc1074.fasta
			cat trimmed-bc1075.for5.fasta trimmed-bc1075.for3.fasta > bc1075.fasta
			cat trimmed-bc1076.for5.fasta trimmed-bc1076.for3.fasta > bc1076.fasta
			cat trimmed-bc1077.for5.fasta trimmed-bc1077.for3.fasta > bc1077.fasta
			cat trimmed-bc1078.for5.fasta trimmed-bc1078.for3.fasta > bc1078.fasta
			cat trimmed-bc1079.for5.fasta trimmed-bc1079.for3.fasta > bc1079.fasta
			cat trimmed-bc1080.for5.fasta trimmed-bc1080.for3.fasta > bc1080.fasta
			cat trimmed-bc1081.for5.fasta trimmed-bc1081.for3.fasta > bc1081.fasta
			cat trimmed-bc1082.for5.fasta trimmed-bc1082.for3.fasta > bc1082.fasta
			cat trimmed-bc1083.for5.fasta trimmed-bc1083.for3.fasta > bc1083.fasta
			cat trimmed-bc1084.for5.fasta trimmed-bc1084.for3.fasta > bc1084.fasta
			cat trimmed-bc1085.for5.fasta trimmed-bc1085.for3.fasta > bc1085.fasta
			cat trimmed-bc1086.for5.fasta trimmed-bc1086.for3.fasta > bc1086.fasta
			cat trimmed-bc1087.for5.fasta trimmed-bc1087.for3.fasta > bc1087.fasta
			cat trimmed-bc1088.for5.fasta trimmed-bc1088.for3.fasta > bc1088.fasta
			cat trimmed-bc1089.for5.fasta trimmed-bc1089.for3.fasta > bc1089.fasta
			cat trimmed-bc1090.for5.fasta trimmed-bc1090.for3.fasta > bc1090.fasta
			cat trimmed-bc1091.for5.fasta trimmed-bc1091.for3.fasta > bc1091.fasta
			cat trimmed-bc1092.for5.fasta trimmed-bc1092.for3.fasta > bc1092.fasta
			cat trimmed-bc1093.for5.fasta trimmed-bc1093.for3.fasta > bc1093.fasta
			cat trimmed-bc1094.for5.fasta trimmed-bc1094.for3.fasta > bc1094.fasta
			cat trimmed-bc1095.for5.fasta trimmed-bc1095.for3.fasta > bc1095.fasta
			cat trimmed-bc1096.for5.fasta trimmed-bc1096.for3.fasta > bc1096.fasta

			rm *.for5.fasta *.for3.fasta

			#demultiplex based on reverse UMI
			for w in bc*.fasta;do
			    fwdumi="${w%%.*}"
			    cutadapt -j $((cores-1)) -e 2 -O 12 -a file:/home/innovation-admin/PRIMERS/PacBIo_V2_UMI97-192.fasta -o "$fwdumi-{name}.fasta" $w #search for all 96 reverse UMIs at 3' end
			done
			
			#delete files with unknown UMIs and files with 0 bytes
			rm *unknown.fasta
			find -name "*.fasta" -size 0 -delete
		fi

		##### symmetrical UMIs (96 + 96)
		if [[ $umitype == "sym" ]]; then

			echo -e "\n****************** Demultiplexing 96X1 UMIs for $plateid...\n"
			#demultiplex (allow up to 2 mismatches and indels)
			cutadapt -j $((cores-1)) -e 2 -O 12 -g file:/home/innovation-admin/PRIMERS/PacBio_V2_UMI1-96.fasta -o "trimmed-{name}.for5.fasta" all.fasta #search for all 96 UMIs at 5' end
			seqtk seq -r trimmed-unknown.for5.fastq > trimmed-unknown.for5.rc.fasta #rev-comp reads that don't have 5'UMI
			cutadapt -j $((cores-1)) -e 2 -O 12 -g file:/home/innovation-admin/PRIMERS/PacBio_V2_UMI1-96.fasta -o "trimmed-{name}.for3.fasta" trimmed-unknown.for5.rc.fasta #search for all 96 UMIs at new 5' end
			rm *unknown*.fasta all.fasta

			#merge forward and reverse split files into a single fasta file
			cat trimmed-bc1001.for5.fasta trimmed-bc1001.for3.fasta > bc1001-bc1001_rc.fasta
			cat trimmed-bc1002.for5.fasta trimmed-bc1002.for3.fasta > bc1002-bc1002_rc.fasta
			cat trimmed-bc1003.for5.fasta trimmed-bc1003.for3.fasta > bc1003-bc1003_rc.fasta
			cat trimmed-bc1004.for5.fasta trimmed-bc1004.for3.fasta > bc1004-bc1004_rc.fasta
			cat trimmed-bc1005.for5.fasta trimmed-bc1005.for3.fasta > bc1005-bc1005_rc.fasta
			cat trimmed-bc1006.for5.fasta trimmed-bc1006.for3.fasta > bc1006-bc1006_rc.fasta
			cat trimmed-bc1007.for5.fasta trimmed-bc1007.for3.fasta > bc1007-bc1007_rc.fasta
			cat trimmed-bc1008.for5.fasta trimmed-bc1008.for3.fasta > bc1008-bc1008_rc.fasta
			cat trimmed-bc1009.for5.fasta trimmed-bc1009.for3.fasta > bc1009-bc1009_rc.fasta
			cat trimmed-bc1010.for5.fasta trimmed-bc1010.for3.fasta > bc1010-bc1010_rc.fasta
			cat trimmed-bc1011.for5.fasta trimmed-bc1011.for3.fasta > bc1011-bc1011_rc.fasta
			cat trimmed-bc1012.for5.fasta trimmed-bc1012.for3.fasta > bc1012-bc1012_rc.fasta
			cat trimmed-bc1013.for5.fasta trimmed-bc1013.for3.fasta > bc1013-bc1013_rc.fasta
			cat trimmed-bc1014.for5.fasta trimmed-bc1014.for3.fasta > bc1014-bc1014_rc.fasta
			cat trimmed-bc1015.for5.fasta trimmed-bc1015.for3.fasta > bc1015-bc1015_rc.fasta
			cat trimmed-bc1016.for5.fasta trimmed-bc1016.for3.fasta > bc1016-bc1016_rc.fasta
			cat trimmed-bc1017.for5.fasta trimmed-bc1017.for3.fasta > bc1017-bc1017_rc.fasta
			cat trimmed-bc1018.for5.fasta trimmed-bc1018.for3.fasta > bc1018-bc1018_rc.fasta
			cat trimmed-bc1019.for5.fasta trimmed-bc1019.for3.fasta > bc1019-bc1019_rc.fasta
			cat trimmed-bc1020.for5.fasta trimmed-bc1020.for3.fasta > bc1020-bc1020_rc.fasta
			cat trimmed-bc1021.for5.fasta trimmed-bc1021.for3.fasta > bc1021-bc1021_rc.fasta
			cat trimmed-bc1022.for5.fasta trimmed-bc1022.for3.fasta > bc1022-bc1022_rc.fasta
			cat trimmed-bc1023.for5.fasta trimmed-bc1023.for3.fasta > bc1023-bc1023_rc.fasta
			cat trimmed-bc1024.for5.fasta trimmed-bc1024.for3.fasta > bc1024-bc1024_rc.fasta
			cat trimmed-bc1025.for5.fasta trimmed-bc1025.for3.fasta > bc1025-bc1025_rc.fasta
			cat trimmed-bc1026.for5.fasta trimmed-bc1026.for3.fasta > bc1026-bc1026_rc.fasta
			cat trimmed-bc1027.for5.fasta trimmed-bc1027.for3.fasta > bc1027-bc1027_rc.fasta
			cat trimmed-bc1028.for5.fasta trimmed-bc1028.for3.fasta > bc1028-bc1028_rc.fasta
			cat trimmed-bc1029.for5.fasta trimmed-bc1029.for3.fasta > bc1029-bc1029_rc.fasta
			cat trimmed-bc1030.for5.fasta trimmed-bc1030.for3.fasta > bc1030-bc1030_rc.fasta
			cat trimmed-bc1031.for5.fasta trimmed-bc1031.for3.fasta > bc1031-bc1031_rc.fasta
			cat trimmed-bc1032.for5.fasta trimmed-bc1032.for3.fasta > bc1032-bc1032_rc.fasta
			cat trimmed-bc1033.for5.fasta trimmed-bc1033.for3.fasta > bc1033-bc1033_rc.fasta
			cat trimmed-bc1034.for5.fasta trimmed-bc1034.for3.fasta > bc1034-bc1034_rc.fasta
			cat trimmed-bc1035.for5.fasta trimmed-bc1035.for3.fasta > bc1035-bc1035_rc.fasta
			cat trimmed-bc1036.for5.fasta trimmed-bc1036.for3.fasta > bc1036-bc1036_rc.fasta
			cat trimmed-bc1037.for5.fasta trimmed-bc1037.for3.fasta > bc1037-bc1037_rc.fasta
			cat trimmed-bc1038.for5.fasta trimmed-bc1038.for3.fasta > bc1038-bc1038_rc.fasta
			cat trimmed-bc1039.for5.fasta trimmed-bc1039.for3.fasta > bc1039-bc1039_rc.fasta
			cat trimmed-bc1040.for5.fasta trimmed-bc1040.for3.fasta > bc1040-bc1040_rc.fasta
			cat trimmed-bc1041.for5.fasta trimmed-bc1041.for3.fasta > bc1041-bc1041_rc.fasta
			cat trimmed-bc1042.for5.fasta trimmed-bc1042.for3.fasta > bc1042-bc1042_rc.fasta
			cat trimmed-bc1043.for5.fasta trimmed-bc1043.for3.fasta > bc1043-bc1043_rc.fasta
			cat trimmed-bc1044.for5.fasta trimmed-bc1044.for3.fasta > bc1044-bc1044_rc.fasta
			cat trimmed-bc1045.for5.fasta trimmed-bc1045.for3.fasta > bc1045-bc1045_rc.fasta
			cat trimmed-bc1046.for5.fasta trimmed-bc1046.for3.fasta > bc1046-bc1046_rc.fasta
			cat trimmed-bc1047.for5.fasta trimmed-bc1047.for3.fasta > bc1047-bc1047_rc.fasta
			cat trimmed-bc1048.for5.fasta trimmed-bc1048.for3.fasta > bc1048-bc1048_rc.fasta
			cat trimmed-bc1049.for5.fasta trimmed-bc1049.for3.fasta > bc1049-bc1049_rc.fasta
			cat trimmed-bc1050.for5.fasta trimmed-bc1050.for3.fasta > bc1050-bc1050_rc.fasta
			cat trimmed-bc1051.for5.fasta trimmed-bc1051.for3.fasta > bc1051-bc1051_rc.fasta
			cat trimmed-bc1052.for5.fasta trimmed-bc1052.for3.fasta > bc1052-bc1052_rc.fasta
			cat trimmed-bc1053.for5.fasta trimmed-bc1053.for3.fasta > bc1053-bc1053_rc.fasta
			cat trimmed-bc1054.for5.fasta trimmed-bc1054.for3.fasta > bc1054-bc1054_rc.fasta
			cat trimmed-bc1055.for5.fasta trimmed-bc1055.for3.fasta > bc1055-bc1055_rc.fasta
			cat trimmed-bc1056.for5.fasta trimmed-bc1056.for3.fasta > bc1056-bc1056_rc.fasta
			cat trimmed-bc1057.for5.fasta trimmed-bc1057.for3.fasta > bc1057-bc1057_rc.fasta
			cat trimmed-bc1058.for5.fasta trimmed-bc1058.for3.fasta > bc1058-bc1058_rc.fasta
			cat trimmed-bc1059.for5.fasta trimmed-bc1059.for3.fasta > bc1059-bc1059_rc.fasta
			cat trimmed-bc1060.for5.fasta trimmed-bc1060.for3.fasta > bc1060-bc1060_rc.fasta
			cat trimmed-bc1061.for5.fasta trimmed-bc1061.for3.fasta > bc1061-bc1061_rc.fasta
			cat trimmed-bc1062.for5.fasta trimmed-bc1062.for3.fasta > bc1062-bc1062_rc.fasta
			cat trimmed-bc1063.for5.fasta trimmed-bc1063.for3.fasta > bc1063-bc1063_rc.fasta
			cat trimmed-bc1064.for5.fasta trimmed-bc1064.for3.fasta > bc1064-bc1064_rc.fasta
			cat trimmed-bc1065.for5.fasta trimmed-bc1065.for3.fasta > bc1065-bc1065_rc.fasta
			cat trimmed-bc1066.for5.fasta trimmed-bc1066.for3.fasta > bc1066-bc1066_rc.fasta
			cat trimmed-bc1067.for5.fasta trimmed-bc1067.for3.fasta > bc1067-bc1067_rc.fasta
			cat trimmed-bc1068.for5.fasta trimmed-bc1068.for3.fasta > bc1068-bc1068_rc.fasta
			cat trimmed-bc1069.for5.fasta trimmed-bc1069.for3.fasta > bc1069-bc1069_rc.fasta
			cat trimmed-bc1070.for5.fasta trimmed-bc1070.for3.fasta > bc1070-bc1070_rc.fasta
			cat trimmed-bc1071.for5.fasta trimmed-bc1071.for3.fasta > bc1071-bc1071_rc.fasta
			cat trimmed-bc1072.for5.fasta trimmed-bc1072.for3.fasta > bc1072-bc1072_rc.fasta
			cat trimmed-bc1073.for5.fasta trimmed-bc1073.for3.fasta > bc1073-bc1073_rc.fasta
			cat trimmed-bc1074.for5.fasta trimmed-bc1074.for3.fasta > bc1074-bc1074_rc.fasta
			cat trimmed-bc1075.for5.fasta trimmed-bc1075.for3.fasta > bc1075-bc1075_rc.fasta
			cat trimmed-bc1076.for5.fasta trimmed-bc1076.for3.fasta > bc1076-bc1076_rc.fasta
			cat trimmed-bc1077.for5.fasta trimmed-bc1077.for3.fasta > bc1077-bc1077_rc.fasta
			cat trimmed-bc1078.for5.fasta trimmed-bc1078.for3.fasta > bc1078-bc1078_rc.fasta
			cat trimmed-bc1079.for5.fasta trimmed-bc1079.for3.fasta > bc1079-bc1079_rc.fasta
			cat trimmed-bc1080.for5.fasta trimmed-bc1080.for3.fasta > bc1080-bc1080_rc.fasta
			cat trimmed-bc1081.for5.fasta trimmed-bc1081.for3.fasta > bc1081-bc1081_rc.fasta
			cat trimmed-bc1082.for5.fasta trimmed-bc1082.for3.fasta > bc1082-bc1082_rc.fasta
			cat trimmed-bc1083.for5.fasta trimmed-bc1083.for3.fasta > bc1083-bc1083_rc.fasta
			cat trimmed-bc1084.for5.fasta trimmed-bc1084.for3.fasta > bc1084-bc1084_rc.fasta
			cat trimmed-bc1085.for5.fasta trimmed-bc1085.for3.fasta > bc1085-bc1085_rc.fasta
			cat trimmed-bc1086.for5.fasta trimmed-bc1086.for3.fasta > bc1086-bc1086_rc.fasta
			cat trimmed-bc1087.for5.fasta trimmed-bc1087.for3.fasta > bc1087-bc1087_rc.fasta
			cat trimmed-bc1088.for5.fasta trimmed-bc1088.for3.fasta > bc1088-bc1088_rc.fasta
			cat trimmed-bc1089.for5.fasta trimmed-bc1089.for3.fasta > bc1089-bc1089_rc.fasta
			cat trimmed-bc1090.for5.fasta trimmed-bc1090.for3.fasta > bc1090-bc1090_rc.fasta
			cat trimmed-bc1091.for5.fasta trimmed-bc1091.for3.fasta > bc1091-bc1091_rc.fasta
			cat trimmed-bc1092.for5.fasta trimmed-bc1092.for3.fasta > bc1092-bc1092_rc.fasta
			cat trimmed-bc1093.for5.fasta trimmed-bc1093.for3.fasta > bc1093-bc1093_rc.fasta
			cat trimmed-bc1094.for5.fasta trimmed-bc1094.for3.fasta > bc1094-bc1094_rc.fasta
			cat trimmed-bc1095.for5.fasta trimmed-bc1095.for3.fasta > bc1095-bc1095_rc.fasta
			cat trimmed-bc1096.for5.fasta trimmed-bc1096.for3.fasta > bc1096-bc1096_rc.fasta

			rm *.for5.fasta *.for3.fasta
			
			#delete files with 0 bytes
			find -name "*.fasta" -size 0 -delete
		fi

		##### asymmetrical Sequel UMIs (96 x 96)
		if [[ $umitype == "sequel" ]]; then

			echo -e "\n****************** Demultiplexing 96X96 UMIs for $plateid...\n"
			#demultiplex (allow up to 2 mismatches and indels)
			cutadapt -j $((cores-1)) -e 2 -O 12 -g file:/home/innovation-admin/PRIMERS/PacBio_V2_UMI1-96.fasta -o "trimmed-{name}.for5.fasta" all.fasta #search for all 96 UMIs at 5' end
			seqtk seq -r trimmed-unknown.for5.fasta > trimmed-unknown.for5.rc.fasta #rev-comp reads that don't have 5'UMI
			cutadapt -j $((cores-1)) -e 2 -O 12 -g file:/home/innovation-admin/PRIMERS/PacBio_V2_UMI1-96.fasta -o "trimmed-{name}.for3.fasta" trimmed-unknown.for5.rc.fasta #search for all 96 UMIs at new 5' end
			rm *unknown*.fasta all.fasta

			#merge forward and reverse split files into a single fasta file
			cat trimmed-bc1001.for5.fasta trimmed-bc1001.for3.fasta > bc1001.fasta
			cat trimmed-bc1002.for5.fasta trimmed-bc1002.for3.fasta > bc1002.fasta
			cat trimmed-bc1003.for5.fasta trimmed-bc1003.for3.fasta > bc1003.fasta
			cat trimmed-bc1004.for5.fasta trimmed-bc1004.for3.fasta > bc1004.fasta
			cat trimmed-bc1005.for5.fasta trimmed-bc1005.for3.fasta > bc1005.fasta
			cat trimmed-bc1006.for5.fasta trimmed-bc1006.for3.fasta > bc1006.fasta
			cat trimmed-bc1007.for5.fasta trimmed-bc1007.for3.fasta > bc1007.fasta
			cat trimmed-bc1008.for5.fasta trimmed-bc1008.for3.fasta > bc1008.fasta
			cat trimmed-bc1009.for5.fasta trimmed-bc1009.for3.fasta > bc1009.fasta
			cat trimmed-bc1010.for5.fasta trimmed-bc1010.for3.fasta > bc1010.fasta
			cat trimmed-bc1011.for5.fasta trimmed-bc1011.for3.fasta > bc1011.fasta
			cat trimmed-bc1012.for5.fasta trimmed-bc1012.for3.fasta > bc1012.fasta
			cat trimmed-bc1013.for5.fasta trimmed-bc1013.for3.fasta > bc1013.fasta
			cat trimmed-bc1014.for5.fasta trimmed-bc1014.for3.fasta > bc1014.fasta
			cat trimmed-bc1015.for5.fasta trimmed-bc1015.for3.fasta > bc1015.fasta
			cat trimmed-bc1016.for5.fasta trimmed-bc1016.for3.fasta > bc1016.fasta
			cat trimmed-bc1017.for5.fasta trimmed-bc1017.for3.fasta > bc1017.fasta
			cat trimmed-bc1018.for5.fasta trimmed-bc1018.for3.fasta > bc1018.fasta
			cat trimmed-bc1019.for5.fasta trimmed-bc1019.for3.fasta > bc1019.fasta
			cat trimmed-bc1020.for5.fasta trimmed-bc1020.for3.fasta > bc1020.fasta
			cat trimmed-bc1021.for5.fasta trimmed-bc1021.for3.fasta > bc1021.fasta
			cat trimmed-bc1022.for5.fasta trimmed-bc1022.for3.fasta > bc1022.fasta
			cat trimmed-bc1023.for5.fasta trimmed-bc1023.for3.fasta > bc1023.fasta
			cat trimmed-bc1024.for5.fasta trimmed-bc1024.for3.fasta > bc1024.fasta
			cat trimmed-bc1025.for5.fasta trimmed-bc1025.for3.fasta > bc1025.fasta
			cat trimmed-bc1026.for5.fasta trimmed-bc1026.for3.fasta > bc1026.fasta
			cat trimmed-bc1027.for5.fasta trimmed-bc1027.for3.fasta > bc1027.fasta
			cat trimmed-bc1028.for5.fasta trimmed-bc1028.for3.fasta > bc1028.fasta
			cat trimmed-bc1029.for5.fasta trimmed-bc1029.for3.fasta > bc1029.fasta
			cat trimmed-bc1030.for5.fasta trimmed-bc1030.for3.fasta > bc1030.fasta
			cat trimmed-bc1031.for5.fasta trimmed-bc1031.for3.fasta > bc1031.fasta
			cat trimmed-bc1032.for5.fasta trimmed-bc1032.for3.fasta > bc1032.fasta
			cat trimmed-bc1033.for5.fasta trimmed-bc1033.for3.fasta > bc1033.fasta
			cat trimmed-bc1034.for5.fasta trimmed-bc1034.for3.fasta > bc1034.fasta
			cat trimmed-bc1035.for5.fasta trimmed-bc1035.for3.fasta > bc1035.fasta
			cat trimmed-bc1036.for5.fasta trimmed-bc1036.for3.fasta > bc1036.fasta
			cat trimmed-bc1037.for5.fasta trimmed-bc1037.for3.fasta > bc1037.fasta
			cat trimmed-bc1038.for5.fasta trimmed-bc1038.for3.fasta > bc1038.fasta
			cat trimmed-bc1039.for5.fasta trimmed-bc1039.for3.fasta > bc1039.fasta
			cat trimmed-bc1040.for5.fasta trimmed-bc1040.for3.fasta > bc1040.fasta
			cat trimmed-bc1041.for5.fasta trimmed-bc1041.for3.fasta > bc1041.fasta
			cat trimmed-bc1042.for5.fasta trimmed-bc1042.for3.fasta > bc1042.fasta
			cat trimmed-bc1043.for5.fasta trimmed-bc1043.for3.fasta > bc1043.fasta
			cat trimmed-bc1044.for5.fasta trimmed-bc1044.for3.fasta > bc1044.fasta
			cat trimmed-bc1045.for5.fasta trimmed-bc1045.for3.fasta > bc1045.fasta
			cat trimmed-bc1046.for5.fasta trimmed-bc1046.for3.fasta > bc1046.fasta
			cat trimmed-bc1047.for5.fasta trimmed-bc1047.for3.fasta > bc1047.fasta
			cat trimmed-bc1048.for5.fasta trimmed-bc1048.for3.fasta > bc1048.fasta
			cat trimmed-bc1049.for5.fasta trimmed-bc1049.for3.fasta > bc1049.fasta
			cat trimmed-bc1050.for5.fasta trimmed-bc1050.for3.fasta > bc1050.fasta
			cat trimmed-bc1051.for5.fasta trimmed-bc1051.for3.fasta > bc1051.fasta
			cat trimmed-bc1052.for5.fasta trimmed-bc1052.for3.fasta > bc1052.fasta
			cat trimmed-bc1053.for5.fasta trimmed-bc1053.for3.fasta > bc1053.fasta
			cat trimmed-bc1054.for5.fasta trimmed-bc1054.for3.fasta > bc1054.fasta
			cat trimmed-bc1055.for5.fasta trimmed-bc1055.for3.fasta > bc1055.fasta
			cat trimmed-bc1056.for5.fasta trimmed-bc1056.for3.fasta > bc1056.fasta
			cat trimmed-bc1057.for5.fasta trimmed-bc1057.for3.fasta > bc1057.fasta
			cat trimmed-bc1058.for5.fasta trimmed-bc1058.for3.fasta > bc1058.fasta
			cat trimmed-bc1059.for5.fasta trimmed-bc1059.for3.fasta > bc1059.fasta
			cat trimmed-bc1060.for5.fasta trimmed-bc1060.for3.fasta > bc1060.fasta
			cat trimmed-bc1061.for5.fasta trimmed-bc1061.for3.fasta > bc1061.fasta
			cat trimmed-bc1062.for5.fasta trimmed-bc1062.for3.fasta > bc1062.fasta
			cat trimmed-bc1063.for5.fasta trimmed-bc1063.for3.fasta > bc1063.fasta
			cat trimmed-bc1064.for5.fasta trimmed-bc1064.for3.fasta > bc1064.fasta
			cat trimmed-bc1065.for5.fasta trimmed-bc1065.for3.fasta > bc1065.fasta
			cat trimmed-bc1066.for5.fasta trimmed-bc1066.for3.fasta > bc1066.fasta
			cat trimmed-bc1067.for5.fasta trimmed-bc1067.for3.fasta > bc1067.fasta
			cat trimmed-bc1068.for5.fasta trimmed-bc1068.for3.fasta > bc1068.fasta
			cat trimmed-bc1069.for5.fasta trimmed-bc1069.for3.fasta > bc1069.fasta
			cat trimmed-bc1070.for5.fasta trimmed-bc1070.for3.fasta > bc1070.fasta
			cat trimmed-bc1071.for5.fasta trimmed-bc1071.for3.fasta > bc1071.fasta
			cat trimmed-bc1072.for5.fasta trimmed-bc1072.for3.fasta > bc1072.fasta
			cat trimmed-bc1073.for5.fasta trimmed-bc1073.for3.fasta > bc1073.fasta
			cat trimmed-bc1074.for5.fasta trimmed-bc1074.for3.fasta > bc1074.fasta
			cat trimmed-bc1075.for5.fasta trimmed-bc1075.for3.fasta > bc1075.fasta
			cat trimmed-bc1076.for5.fasta trimmed-bc1076.for3.fasta > bc1076.fasta
			cat trimmed-bc1077.for5.fasta trimmed-bc1077.for3.fasta > bc1077.fasta
			cat trimmed-bc1078.for5.fasta trimmed-bc1078.for3.fasta > bc1078.fasta
			cat trimmed-bc1079.for5.fasta trimmed-bc1079.for3.fasta > bc1079.fasta
			cat trimmed-bc1080.for5.fasta trimmed-bc1080.for3.fasta > bc1080.fasta
			cat trimmed-bc1081.for5.fasta trimmed-bc1081.for3.fasta > bc1081.fasta
			cat trimmed-bc1082.for5.fasta trimmed-bc1082.for3.fasta > bc1082.fasta
			cat trimmed-bc1083.for5.fasta trimmed-bc1083.for3.fasta > bc1083.fasta
			cat trimmed-bc1084.for5.fasta trimmed-bc1084.for3.fasta > bc1084.fasta
			cat trimmed-bc1085.for5.fasta trimmed-bc1085.for3.fasta > bc1085.fasta
			cat trimmed-bc1086.for5.fasta trimmed-bc1086.for3.fasta > bc1086.fasta
			cat trimmed-bc1087.for5.fasta trimmed-bc1087.for3.fasta > bc1087.fasta
			cat trimmed-bc1088.for5.fasta trimmed-bc1088.for3.fasta > bc1088.fasta
			cat trimmed-bc1089.for5.fasta trimmed-bc1089.for3.fasta > bc1089.fasta
			cat trimmed-bc1090.for5.fasta trimmed-bc1090.for3.fasta > bc1090.fasta
			cat trimmed-bc1091.for5.fasta trimmed-bc1091.for3.fasta > bc1091.fasta
			cat trimmed-bc1092.for5.fasta trimmed-bc1092.for3.fasta > bc1092.fasta
			cat trimmed-bc1093.for5.fasta trimmed-bc1093.for3.fasta > bc1093.fasta
			cat trimmed-bc1094.for5.fasta trimmed-bc1094.for3.fasta > bc1094.fasta
			cat trimmed-bc1095.for5.fasta trimmed-bc1095.for3.fasta > bc1095.fasta
			cat trimmed-bc1096.for5.fasta trimmed-bc1096.for3.fasta > bc1096.fasta

			rm *.for5.fasta *.for3.fasta

			#demultiplex based on reverse UMI
			for w in bc*.fasta;do
			    fwdumi="${w%%.*}"
			    cutadapt -j $((cores-1)) -e 2 -O 12 -a file:/home/innovation-admin/PRIMERS/PacBIo_V2_UMI97-192_SQL.fasta -o "$fwdumi-{name}.fasta" $w #search for all 96 reverse UMIs at 3' end
			done

			#delete files with unknown UMIs and files with 0 bytes
			rm *unknown.fasta
			find -name "*.fasta" -size 0 -delete
		fi

		#rename files based on UMI mapping file
		while IFS=$'\t' read -r fwd rev samp group umiplateid sampleplateid; do
		    old_name="${fwd}-${rev}.fasta"
		    old_name=$(echo "$old_name" | tr -d '[:space:]')
		    new_name="${samp}.fasta"
		    mv "/home/innovation-admin/DATA_INPUT/"$plateid"/$old_name" "/home/innovation-admin/DATA_INPUT/"$plateid"/$new_name"
		done < /home/innovation-admin/DATA_INPUT/mapping_"$plateid".txt

		#delete files with 0 bytes
		find -name "*.fasta" -size 0 -delete

		#delete files with UMI combinations that were not renamed due to omission from UMI mapping file
		rm bc*.fasta

		#filter by read size again
		for r in *.fasta;do cutadapt -j $((cores-1)) -m $minreadlen -M $maxreadlen -o $r.filt $r;done
		rm *.fasta
		rename 's/.filt//g' *.filt

		#delete files with 0 bytes
		find -name "*.fasta" -size 0 -delete

		#get read count of demultiplexed reads
		sum=0
		for k in *.fasta;do sum=$((sum + $(echo $(cat $k|wc -l)/2+1|bc)));done
		umidemuxreadcount=$sum
	fi

	#if UMI demultiplexing is the last step, move to the next plate (if exists), otherwise skip to the end of the plate loop
	if [[ $umidemuxstep == "yes" && $ptrimstep == "no" && $contigstep == "no" ]]; then
		return
	fi

	
# <<<<<<<<<<< BY THIS POINT, ALL DATA SHOULD BE INDIVIDUAL FASTA FILES NAMED WITH SAMPLE IDS


	sampletask(){
	    #get sample name
	    sampleid="${i%.*}"

	    ########################################################################################################################################################################
		# primer trimming
		########################################################################################################################################################################

		#if primer trim not requested, rename sequence headers to shorter version
		if [[ $ptrimstep == "no" ]]; then
			#rename trimmed read headers to shorter versions
		    cut -d" " -f1 "$sampleid".fasta > "$sampleid".fas
		    rm "$sampleid".fasta
		    mv "$sampleid".fas "$sampleid".fasta		    
		fi

		#if primer trim requested, proceed with trimming
		if [[ $ptrimstep == "yes" ]]; then
		    echo -e "******** Trimming primers from $sampleid reads..."

		    #trim primers
		    cutadapt -j 1 -e 0.2 -O 12 -g file:temp_fwd_primers.fa --untrimmed-output "$sampleid"_fwd5_noprimer.fa -o "$sampleid"_fwd5_withprimer.fa $i #search forward primer at 5' end
		    cutadapt -j 1 -e 0.2 -O 12 -a file:temp_rev_primers_rc.fa --discard-untrimmed -o "$sampleid"_rev3_withprimer.fa "$sampleid"_fwd5_withprimer.fa #trim reverse primer from 3' end
		    cutadapt -j 1 -e 0.2 -O 12 -g file:temp_rev_primers.fa --discard-untrimmed -o "$sampleid"_rev5_withprimer.fa "$sampleid"_fwd5_noprimer.fa #search reverse primer at 5' end of reads with no fwd primer at 5' end
		    cutadapt -j 1 -e 0.2 -O 12 -a file:temp_fwd_primers_rc.fa --discard-untrimmed -o "$sampleid"_fwd3_withprimer.fa "$sampleid"_rev5_withprimer.fa #trim forward primer from 3' end

		    #replace empty reads with N and rev-comp backwards reads
		    sed '2~4s/^$/N/; 4~4s/^$/S/' "$sampleid"_fwd3_withprimer.fa > "$sampleid"_fwd3_withprimer_noempty.fa
		    seqtk seq -r "$sampleid"_fwd3_withprimer_noempty.fa > "$sampleid"_fwd3_withprimer.rc.fa

		    #merge all reads into a single FASTQ file
		    cat "$sampleid"_rev3_withprimer.fa "$sampleid"_fwd3_withprimer.rc.fa > "$sampleid".fas
		    
		    #delete intermediate files
		    rm "$i" "$sampleid"_fwd5_noprimer.fa "$sampleid"_fwd5_withprimer.fa "$sampleid"_rev5_withprimer.fa
		    rm "$sampleid"_fwd3_withprimer.fa "$sampleid"_fwd3_withprimer.rc.fa "$sampleid"_rev3_withprimer.fa "$sampleid"_fwd3_withprimer_noempty.fa
		    
		    #size filter again
		    echo -e "******** Size filtering post-primer-trimmed $sampleid reads..."
		    cutadapt -j 1 -m $minreadlen -M $maxreadlen -o "$sampleid".fasta "$sampleid".fas
		    rm "$sampleid".fas

		    #rename FASTA file to _trimmed.fasta
		    rename 's/.fasta/_trimmed.fasta/g' "$sampleid".fasta
		    
		    #rename trimmed read headers to shorter versions
		    cut -d" " -f1 "$sampleid"_trimmed.fasta > "$sampleid"_trimmed.fas
		    rm "$sampleid"_trimmed.fasta
		    mv "$sampleid"_trimmed.fas "$sampleid".fasta
		fi
	    
	    #if primer trimming is the last step, move to the next sampleID
		if [[ $ptrimstep == "yes" && $contigstep == "no" ]]; then
			return
		fi

	    ########################################################################################################################################################################
		# contig clustering
		########################################################################################################################################################################
		
		#if script has not already exited by this point, contig clustering was requested and no conditional IF statement is necessary
	    
	    #temporarily trim 15% of bases from the ends of each read
	    echo -e "******** Temporarily trimming 15% of bases from ends of $sampleid reads..."
	    cutadapt -j 1 -u $trimlength -o "$sampleid"_trimmed.fast "$sampleid".fasta
	    cutadapt -j 1 -u $((-$trimlength)) -o "$sampleid"_trimmed.fas "$sampleid"_trimmed.fast
	    rm "$sampleid"_trimmed.fast

	    #cluster with VSEARCH based on internal portion of reads only
	    echo -e "******** Performing primary clustering of $sampleid..."
	    vsearch --cluster_fast "$sampleid"_trimmed.fas \
	    --id 0.95 \
	    --clusters "$sampleid"_Contig \
	    --iddef 3 \
	    --threads 1

	    #delete temporary trimmed files
	    rm "$sampleid"_trimmed.fas

	    #delete contigs with fewer than minreads
	    echo -e "******** Deleting $sampleid primary contigs with fewer than minreads..."
	    find . -type f -name "$sampleid"_Contig\* | while IFS= read -r file; do
	        read_count=$(grep -c "^>" "$file")
	        if [ "$read_count" -lt $minreads ]; then
	            rm "$file"
	        fi
	    done

	    #rename contig read files to _Reads.fas
	    echo -e "******** Renaming primmary contigs of $sampleid..."
	    find . -type f -name "$sampleid"_Contig\* | while IFS= read -r file; do
	        new_name="${file}_Reads.fas"
	        mv "$file" "$new_name"
	        echo "Renamed: $file -> $new_name"
	    done
	            
	    #Extract read names from each contig file, then pull those reads from the original fasta file to make contigs with full-length reads
	    echo -e "******** Restoring full length reads into $sampleid primary contigs..."
	    for f in "$sampleid"_Contig*_Reads.fas; do
	       grep '^>' "$f" | cut -d '>' -f 2 | grep -A 1 -Ff - "${sampleid}.fasta" | grep -v '^--$' > "$f.realcontig"
	       rm $f
	       rename 's/.fas.realcontig/.fasta/g' "$f.realcontig"
	    done
 	 
	    #generate contig consensus sequences in R
	    echo -e "******** Generating $sampleid primary contig consensus sequences..."
	    Rscript /home/innovation-admin/SCRIPTS/SUBSCRIPTS/primary_contig_consensus.R "$plateid" "$sampleid"

	    #cluster contig consensus sequences
	    echo -e "******** Performing secondary clustering of $sampleid..."
	    vsearch --cluster_fast "$sampleid"_contigs.tmp \
	    --id 0.98 \
	    --clusters "$sampleid"_FinalContig \
	    --iddef 3 \
	    --threads 1

	    #delete original contigs
	    echo -e "******** Deleting $sampleid primary contig consensus sequences..."
	    rm "$sampleid"_contigs.tmp

	    #rename contig read files to _Reads.fas
	    echo -e "******** Renaming secondary contigs of $sampleid..."
	    find . -type f -name "$sampleid"_FinalContig\* | while IFS= read -r file; do
	        new_name="${file}_Reads.fasta"
	        mv "$file" "$new_name"
	        echo "Renamed: $file -> $new_name"
	    done

	    #generate new contig consensus sequences in R
	    echo -e "******** Generating $sampleid secondary contig consensus sequences..."
	    Rscript /home/innovation-admin/SCRIPTS/SUBSCRIPTS/final_contig_consensus.R "$plateid" "$sampleid"

	    #combine original reads into a single file to reflect final contig reads
	    echo -e "******** Restoring orignal reads into $sampleid secondary contigs..."
	    for f in "$sampleid"_FinalContig*_Reads.fasta; do
	        output=$(grep '^>' "$f" | cut -d '>' -f 2 | cut -d '|' -f 1,2 | tr '|' '_')
	        while IFS= read -r line; do
	          cat "$line"_Reads.fasta >> $f.out
	        done <<< "$output"
	        rm $f
	        mv $f.out $f
	    done

	    echo -e "******** Deleting $sampleid primary contigs..."
	    rm "$sampleid"_Contig*_Reads.fasta
	    
	    echo -e "******** Renaming $sampleid secondary contig read files..."
	    rename 's/FinalContig/Contig/g' "$sampleid"_FinalContig*_Reads.fasta
	}

	for i in *.fasta; do
	    sampletask "$i" &
	    if [[ $(jobs -r -p | wc -l) -ge $((cores-2)) ]]; then
	        wait -n
	    fi
	done
	wait

	#delete temporary primer fasta files
	echo -e "******** Deleting temporary primer sequence files for $plateid..."
	rm temp_fwd_primers.fa temp_fwd_primers_rc.fa temp_rev_primers.fa temp_rev_primers_rc.fa

	#if primer trimming was the last step, move to the next plate (if exists) or skip to the end of the plate loop
	if [[ $ptrimstep == "yes" && $contigstep == "no" ]]; then
		echo -e "******** Deleting temporary primer sequence files for $plateid..."
		rm temp_fwd_primers.fa temp_fwd_primers_rc.fa temp_rev_primers.fa temp_rev_primers_rc.fa
		return
	fi

	#delete any original read, contig component read, or contig consensus read files that are empty
	find ./ -name "*.fasta" -size 0 -delete
	find ./ -name "*.tmp" -size 0 -delete

	#Move original FASTA files to their own folder
	mv /home/innovation-admin/DATA_INPUT/"$plateid"/*.fasta /home/innovation-admin/DATA_INPUT/"$plateid"/Original_Reads
	mv /home/innovation-admin/DATA_INPUT/"$plateid"/Original_Reads/*Reads.fasta /home/innovation-admin/DATA_INPUT/"$plateid"/

	#count primer trimmed reads if applicable
	if [[ $ptrimstep == "yes" ]]; then
		echo -e "******** Counting primer trimmed reads..."
		sum=0
		for k in ./Original_Reads/*.fasta;do count=$(grep -c "^>" "$k"); sum=$((sum + count));done
		ptrimreadcount=$sum
	fi

	#move contig read files into their own folder
	echo *Reads.fasta | xargs mv -t Contig_Component_Reads --

	#count contig reads
	echo -e "******** Counting contig reads..."
	sum=0
	for k in ./Contig_Component_Reads/*.fasta;do count=$(grep -c "^>" "$k"); sum=$((sum + count));done
	contigreadcount=$sum

	#rename contig files
	rename 's/tmp/fasta/g' /home/innovation-admin/DATA_INPUT/"$plateid"/*finalcontigs.tmp
	rename 's/finalcontigs/Contigs/g' /home/innovation-admin/DATA_INPUT/"$plateid"/*finalcontigs.fasta

	#merge all contigs into a single file
	echo -e "******** Merging all contig consensus sequences into a single FASTA file..."
	cat /home/innovation-admin/DATA_INPUT/"$plateid"/*Contigs.fasta > /home/innovation-admin/DATA_INPUT/"$plateid"/"$plateid".fasta

	#assign taxonomy to contigs and output summary table
	echo -e "******** Assigning taxonomy to contig consensus sequences..."
	vsearch --sintax /home/innovation-admin/DATA_INPUT/"$plateid"/"$plateid".fasta \
	-db /home/innovation-admin/REFS/$reflib.fasta \
	-tabbedout /home/innovation-admin/DATA_INPUT/"$plateid"/"$plateid".table \
	-strand plus \
	-sintax_cutoff $sintax \
	-threads $((cores-1))

	#reformat tax id table
	echo -e "******** Reformatting SINTAX ID table..."
	awk 'BEGIN{OFS="\t"; print "SeqName","TaxAssign","Strand","TaxAssignFinal";}{print}' "$plateid".table > tmpfile && mv tmpfile "$plateid".table

	#add contig sequences to tax ID table, filter by taxonomy (if supplied), auto-trim sequences, and extract dominant contigs
	echo -e "******** Auto-trim, extracting dominant contigs, etc..."
	Rscript /home/innovation-admin/SCRIPTS/SUBSCRIPTS/autotrim_taxfilter_dominants.R "$plateid" "$boldtaxpresent"
	rm "$plateid".table
	rm "$plateid".fasta #delete reads that were not auto-trimmed

	#count reads in dominant contigs
	echo -e "******** Counting reads in dominant contigs..."
	if grep ">" /home/innovation-admin/DATA_INPUT/"$plateid"/"$plateid"_DominantContigs.fasta; then
			dominantreadcount=$(grep ">" /home/innovation-admin/DATA_INPUT/"$plateid"/"$plateid"_DominantContigs.fasta | cut -d'|' -f3 | cut -d'-' -f2 | paste -sd+ - | bc)
	fi

	#generate reaad count text file for plate/sample
	echo -e "Input reads:$inputreadcount" > "$plateid"_readcounts.txt
	echo -e "After basecalling:$basecallreadcount" >> "$plateid"_readcounts.txt
	echo -e "After NB demux:$nbdemuxreadcount" >> "$plateid"_readcounts.txt
	echo -e "Assigned to plate/sample:$plateassignreadcount" >> "$plateid"_readcounts.txt
	echo -e "After size filtering:$sizefilterreadcount" >> "$plateid"_readcounts.txt
	echo -e "After UMI demux:$umidemuxreadcount" >> "$plateid"_readcounts.txt
	echo -e "After primer trim:$ptrimreadcount" >> "$plateid"_readcounts.txt
	echo -e "Reads in contigs:$contigreadcount" >> "$plateid"_readcounts.txt
	echo -e "Reads in dominants:$dominantreadcount" >> "$plateid"_readcounts.txt
	
	#generate summary histograms for plate/sample
	echo -e "******** Generating histograms..."
	Rscript /home/innovation-admin/SCRIPTS/SUBSCRIPTS/generate_histograms.R "$plateid" "$nbdemuxstep"

	#move contig files into folder
	mv /home/innovation-admin/DATA_INPUT/"$plateid"/*Contigs.fasta /home/innovation-admin/DATA_INPUT/"$plateid"/Contig_Consensus_Sequences
	mv /home/innovation-admin/DATA_INPUT/"$plateid"/Contig_Consensus_Sequences/"$plateid"_AllContigs.fasta /home/innovation-admin/DATA_INPUT/"$plateid"/
	mv /home/innovation-admin/DATA_INPUT/"$plateid"/Contig_Consensus_Sequences/"$plateid"_DominantContigs.fasta /home/innovation-admin/DATA_INPUT/"$plateid"/
}

#run the above platetask() on each plate/library folder
for dir in /home/innovation-admin/DATA_INPUT/*/; do 
  platetask "$dir"
done

#if size filtering is the last step, exit
if [[ $sizefilterstep == "yes" && $umidemuxstep == "no" && $ptrimstep == "no" && $contigstep == "no" ]]; then
	echo -e "\nSize filtering was the last step requested. Exiting script."
	exit
fi

#if UMI demultiplexing was the last step, exit
if [[ $umidemuxstep == "yes" && $ptrimstep == "no" && $contigstep == "no" ]]; then
	echo -e "\nUMI demultiplexing was the last step. Exiting script."
	exit
fi

#if primer trimming was the last step, exit
if [[ $ptrimstep == "yes" && $contigstep == "no" ]]; then
	echo -e "\nPrimer trimming was the last step requested. Exiting script."
	exit
fi


#if script makes it to this point, contig assembly was the last step, so package everything up
cd /home/innovation-admin/DATA_INPUT/

mkdir -m 777 METADATA_FILES
mv *.txt *.xlsx METADATA_FILES

mkdir -m 777 /home/innovation-admin/DATA_INPUT/"$runid"
mv /home/innovation-admin/DATA_INPUT/* /home/innovation-admin/DATA_INPUT/"$runid"

mv -f "$runid" /home/innovation-admin/DATA_OUTPUT

	
```
