## Supplemental File 1 for "Barcode 100K Specimens: In a Single Nanopore Run": pairwise_divergence.py

```
#!/usr/bin/env python3
# -*- coding: utf-8 -*-
"""
Created on Fri Jun 16 16:22:27 2023
Takes a fasta file containing pairs of sequences to be compared and outputs
a table of pairwise divergence % values. The specimen name is taken as any text
up to the first "|" character in the fasta header. Optionally, sequences may
have a suffix following a "." character which will also be ignored by this
script and only the text before the "." will be treated as the name of the 
specimen for which to compare a pair of sequences. 
Only generates results where exactly two sequences per specimen name are found.
If a specimen has only one sequence, that is ignored and left out of the output.
If a specimen has more than two sequences, that specimen is also excluded from 
comparison and an error message is returned.
Only A/G/C/T are counted as valid bases. Gaps, Ns or other IUPAC codes in either
position being compared mean that position is treated as missing data and ignored.
@author: robinfloyd
"""

import sys
from Bio import Align, SeqIO

def pairwise_align(seq1, seq2): #function to align a pair of sequences   
    alignments = aligner.align(seq1, seq2)    
    alignment = alignments[0]    
    aln1, aln2 = alignment
    return aln1, aln2

def calculate_divergence(aln1,aln2): #function to calculate divergence from an aligned pair
    mismatches = 0
    comparison_length=0
    for i in range(0,len(aln1)):
        aln1_base = aln1[i]
        aln2_base = aln2[i]
                        
        if (aln1_base in valid_bases) and (aln2_base in valid_bases):  
            comparison_length += 1
            if aln1_base != aln2_base:
                mismatches += 1
    divergence = 100*(mismatches/comparison_length)
    return divergence

infile = sys.argv[1]
outfilename = infile.rsplit('.')[0] + '_divergence_table.csv'

aligner = Align.PairwiseAligner() # Alignment module from Biopython. Parameters set below
aligner.match_score = 1.0 
aligner.open_gap_score = -10
aligner.extend_gap_score = -1

comparison_dict = {}
valid_bases = ['A','G','C','T'] # Only A/G/C/T are counted as valid bases. 
divergence_results ={}

for seq_record in SeqIO.parse(infile, "fasta"): 
    newheader = seq_record.id.split('|')[0]
    # capture any text up to the first pipe character, this is used as the unique 
    # specimen name for which we will compare a pair of sequences
    
    specimenname = newheader.rsplit('.')[0]
    # if any dot character is part of the name, take only the text before the 
    # final dot to use as the specimen name for comparing a pair of sequences
    
    if not (specimenname in comparison_dict.keys()):
        comparison_dict[specimenname] = []
    # if we do not have any sequences from this specimen, create a dictionary
    # entry for the specimen as an empty list
            
    comparison_dict[specimenname].append(seq_record.seq)
    # append the current sequence to the dictionary entry for the specimen

for specimen, seq_pair in comparison_dict.items():
    
    if len(seq_pair) > 2: #if we found more than 2 seqs per specimen, raise an error
        print('ERROR:',specimen,'- more than two sequences found. This tool compares exactly two sequences per specimen.')
        pass
    
    elif len(seq_pair) < 2: #if a specimen has only 1 seq, ignore and move on to the next
        pass
    
    else: #only do the comparison if we have exactly 2 seqs per specimen
        seq1, seq2 = seq_pair
        aln1, aln2 = pairwise_align(seq1, seq2)
        if len(aln1) == len(aln2):       
            divergence_results[specimen] = calculate_divergence(aln1, aln2)           
        else:
            print(specimen,'- alignment error.')
            pass
       
with open(outfilename, 'w') as outfile:
    for specimen, divergence in divergence_results.items():
        outfile.write("%s,%s\n"%(specimen,divergence))
```
