## Supplemental File 1 for "Barcode 100K Specimens: In a Single Nanopore Run": primary_contig_consensus.R

```
#get argument from bash script (plate name)
args <- commandArgs(trailingOnly = TRUE)
plateid <- args[1]
sampleid <- args[2]

if (length(plateid) == 0) {
  stop("Error: Missing input argument (plateid).\n", call. = FALSE)
}

if (length(sampleid) == 0) {
  stop("Error: Missing input argument (sampleid).\n", call. = FALSE)
}

#set working directory based on plate's folder, load required libraries
setwd(paste("/home/innovation-admin/DATA_INPUT/",plateid,"/", sep = ""))
library("Biostrings")
library(muscle)
library(DECIPHER)

#for each contig file
output <- DNAStringSet()

pattern <- paste0(sampleid, "_Contig.*_Reads.fasta")
file_list <- grep(pattern, list.files(), value = TRUE)

for (file in file_list) {
  contigname <- gsub("_Reads.fasta", "", file)
  contigname <- gsub("_Contig", "|Contig", contigname)
  input <- readDNAStringSet(file)
  len <- length(input)
  if (len >= 2) {
    if (len > 1000) {
      indices <- sample(1:(max(len)), 1000, replace=FALSE)
      input <- input[indices]
    }
    align <- DNAStringSet(muscle(input))
    cons <- DNAStringSet(ConsensusSequence(align, ignoreNonBases = FALSE, noConsensusChar = "N",threshold = 0.75, minInformation = 0.25))
    names(cons) <- sprintf("%s|reads-%s", contigname, len)
    #temp <- DNAStringSet(muscle(c(input,cons))) #THIS IS FOR TESTING ONLY
    output <- c(output,cons)
  }
}

#remove all alignment gaps from consensus sequences
output <- DNAStringSet(gsub(pattern = "-", replacement = "", x = output))

#replace any degenerate bases with N
degen <- paste0("[^", paste(c("A","T","C","G","N"), collapse = ""), "]")
output <- DNAStringSet(gsub(pattern = degen, "N", output))

#output consensus sequences as a single FASTA file
writeXStringSet(output,sprintf("%s_contigs.tmp",sampleid))
```
