## Supplemental File 1 for "Barcode 100K Specimens: In a Single Nanopore Run": README.txt

The following scripts are used for all bioinformatic processes described in the manuscript "BARCODE 100K SPECIMENS: IN A SINGLE NANOPORE RUN" by Hebert et al, 2023 (BioRxiv).
The following working directory structure is recommended. The code for each script is currently set to a directory structure used in our lab, and must be modified to reflect your specific folder names and directory structure.

DATA_INPUT - This is the main working directory in which all input data is placed before triggering the master script (ONT.sh)
DATA_OUTPUT - This is the output folder in which completed runs can be found.
SCRIPTS - This folder contains the master script (ONT.sh).
SCRIPTS > SUBSCRIPTS - This subfolder within SCRIPTS contains all dependency R scripts.
REF - This fodler contains all SINTAX formatted reference libraries (FASTA files).
PRIMERS - This folder should contian a FASTA file called PrimersDB.fasta, which contains all forward and reverse primer sequences in 5' -> 3' orientation.

The following lists all scripts with a description of their use:

1) ONT.sh - This is the master script that is called via 'bash ONT.sh'. It requires two input data and one optional file:
	
	i) 	 parameters_XXXXX.xlsx (REQUIRED) - This file allows the user to set analysis parameters, specify primers, specify reference libraries, and provide a UMI map. Detailed instructions are inluded within.
	ii)  Raw data (REQUIRED) - This can be one one of three formats: POD5 file(s) from the sequencer, FASTQ file(s), or FASTA file(s). 
	iii) taxonomy_XXXXX.xlsx (OPTIONAL) - This file can be downloaded from BOLD by searching for all specimens in your sequencing run and downloading the corresponding BOLD datasheet. Be sure to include the defaut lab progress tab as well as the Toxonomy tab.

	All input file should be placed into the working directory, at which point this script can be called from the termianl via 'bash ONT.sh'. The user will then follow the prompts. All dependency scripts are listed below:

2) get_parameters.R - This is a dependency R script used by ONT.sh. It extracts relevent run infomrmation from the input parameters file.

3) primary_contig_consensus.R - This is a dependency R script used by ONT.sh. It generates consensus sequences for the primary contigs.

4) final_contig_consensus.R - This is a dependency R script used by ONT.sh. It generates consensus sequences for the final contings.

5) autotrim_taxfilter_dominants.R - This is a dependency R script used by ONT.sh. It trims the final contig sequences to the precise barcode region, assigns taxonomy, and selects dominant contigs.

6) generate_histograms.R - This is a dependency R script used by ONT.sh. It generates summary histograms that can be used to quickly assess the quality of the run.

The output of ONT.sh is a single folder named after the user-specified run name. Within are the following files/folders:

i)	 METADATA_FILES - This folder contains the input parameters and taxonopmy (if provided) files, as well as the corresponding text files generated by ONT.sh

ii)  terminalhistory.log - This is a text file that contains all terminal standard output and standard errors. It is used for troublshooting if necessary.

iii) LibaryName - This folder contains most of the output data, and will be named after the user-specified library/plate name. If native barcodes were used, there will be a separate folder for each library/plate. The follwing data will be found within each LibraryName folder:

	a)	all.fastq - This is the raw input data resulting from base calling. It is useful if the user wants to repeat analsyis without having to repeat base calling.
	b)	LibraryName_AllContigs.fasta - This FASTA file contains all contig consensus sequences.
	c)	LibraryName_DominantContigs.fasta - This FASTA file contains all dominant contig consensus sequences.
	d)	LibraryName_TaxonomicAssignments_AllContigs.tsv - This text file (can be opened in Excel) is composed of one row for each contig, and contains a lot of information about the contigs including read count, length, taxonomic ID, etc.
	e)	LibraryName_TaxonomicAssignments_DominantContigs.tsv - This text file is the same as the above, but contains only dominant contigs as opposed to all contigs.
	f)	LibraryName_SummaryHistograms - This PDF contains histograms useful for assessing the quality and performance of the run.
	g)	LibraryName_readcounts.txt - This text file was produced by ONT.sh and was used to generate one of the above histograms. It lists read counts after each step in the bioinformatic process.
	h) 	Contig_Component_Reads - This folder contains one FASTA file for each contig, which comprise all raw reads that make up that contig. They are useful for assessing the composition of each contig.
	i)	Contig_Consensus_Sequences - This folder contains one FASTA file for each sample that produced at least one contig. Each file contains the consensus sequence(s) of each contig for that sample.
	j)	Original_Reads - This folder contains one FASTA file for each sample that yielded at least one read, and comprise all pre-clustering reads associated with that sample. They are useful for seeing if samples that failed to yield a contig still had a few reads that were below the user-specified minimum threshold.

The python script below is not involved in the main bioinformatc pipeline, but was used for this study to assess seqeunce congruence between ONT and Sequel-generated barcode sequences:

pairewise_divergence.py - This python script takes as input a single fasta file containing pairs of sequences to be compared. The specimen name is taken as any text up to the first "|" character in the fasta header. Optionally, sequences may have a suffix following a "." character, and only the text before the "." will be treated as the name of the specimen. It aligns each read pair and then compares the nucleotides at each position, counting any discordances. The number of discordances are summed and compared to the original sequence lengths to calculate a percent pairwise divergence. This is done for each sequence pair and the output is a table of divergence values for each sample pair.
